# Alternative LC-MS/MS Platforms and Data Acquisition Strategies for Proteomic Genotyping of Human Hair Shafts

**DOI:** 10.1101/2021.03.15.435505

**Authors:** Zachary C. Goecker, Kevin M. Legg, Michelle R. Salemi, Anthony W. Herren, Brett S. Phinney, Heather E. McKiernan, Glendon J. Parker

## Abstract

Protein is a major component of all biological evidence. Proteomic genotyping is the use of genetically variant peptides that contain single amino acid polymorphisms to infer the genotype of matching non-synonymous single nucleotide polymorphisms for the individual who originated the protein sample. This can be used to statistically associate an individual to evidence found at a crime scene. The utility of the inferred genotype increases as the detection of genetically variant peptides increases, which is the direct result of technology transfer to mass spectrometry platforms typically available. Digests of single (2 cm) human hair shafts from three European and two African subjects were analyzed using data dependent acquisition on a Q-Exactive™ Plus Hybrid Quadrupole-Orbitrap™ system, data independent acquisition and a variant of parallel reaction monitoring on a Orbitrap Fusion™ Lumos™ Tribrid™ system, and multiple reaction monitoring on an Agilent 6495 triple quadrupole system. In our hands, average genetically variant peptide detection from a selected 24 genetically variant peptide panel increased from 6.5 ± 1.1 and 3.1 ± 0.8 using data dependent and independent acquisition to 9.5 ± 0.7 and 11.7 ± 1.7 using parallel reaction monitoring and multiple reaction monitoring (p < 0.05). Parallel reaction monitoring resulted in a 1.3-fold increase in detection sensitivity, and multiple reaction monitoring resulted in a 1.6-fold increase in detection sensitivity. This increase in biomarker detection has a functional impact on the statistical association of a protein sample and an individual. Increased biomarker sensitivity, using Markov Chain Monte Carlo modeling, produced a median estimated random match probability of over 1 in 10 trillion from a single hair using targeted proteomics. For parallel reaction monitoring and multiple reaction monitoring, detected genetically variant peptides were validated by the inclusion of stable isotope labeled peptides in each sample, which served also as a detection trigger. This research accomplishes two aims: the demonstration of utility for alternative analytical platforms in proteomic genotyping, and the establishment of validation methods for the evaluation of inferred genotypes.

**Highlights:**

- Test four mass spectrometry configurations to optimize detection of genetically variant peptides
- Technology transfer of proteomic genotyping assays
- Improved sensitivity results in higher level of forensic discrimination for human identification using multiple reaction monitoring

**Graphical Abstract:** 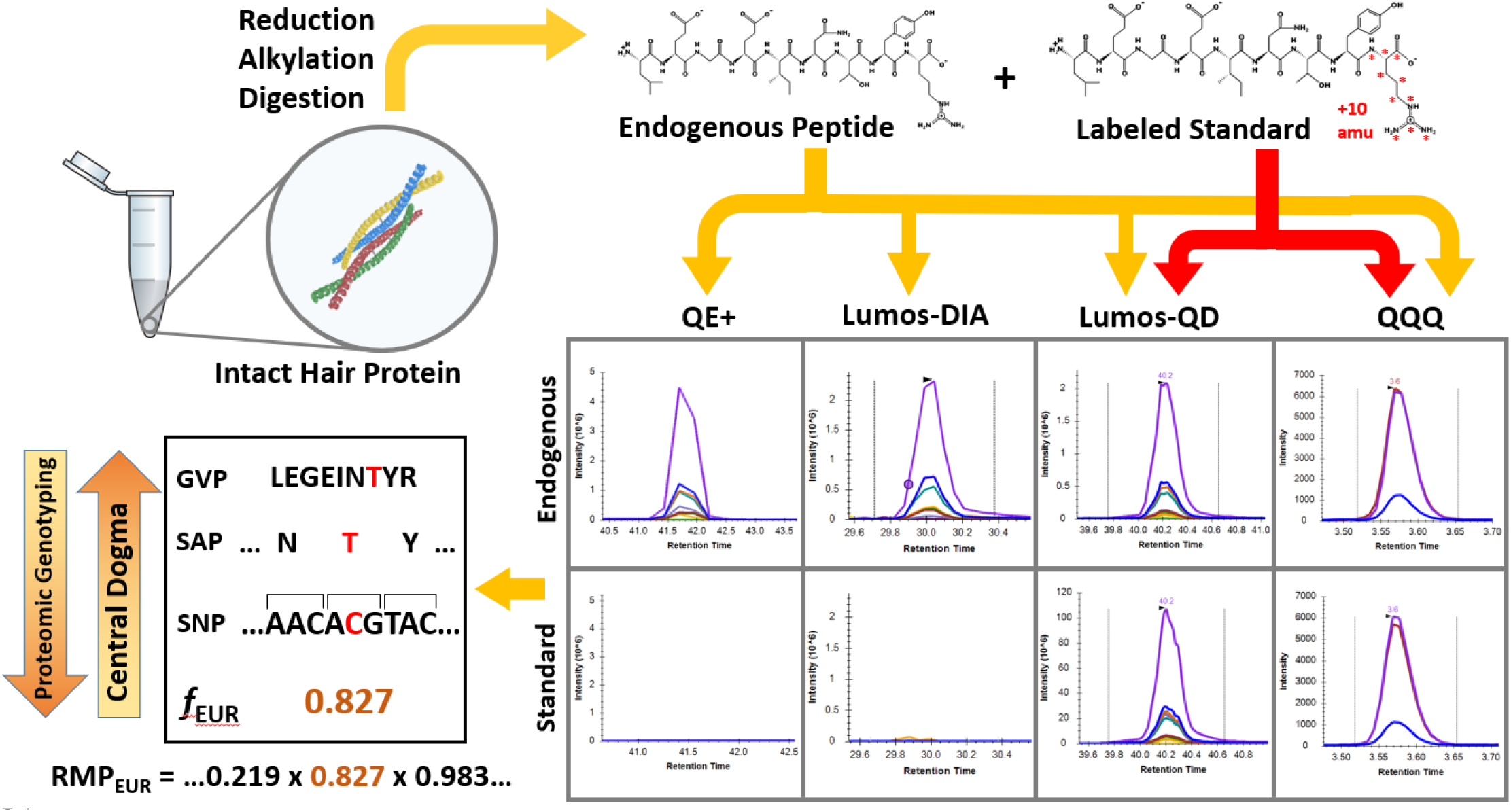

## 1. Introduction

Proteomics has many promising applications in a legal context, with recent advances being made in body fluid identification, drug interactions, sex estimation, and human identification (HID) [1–6]. Proteomic genotyping is the analysis of protein sequence variants, termed single amino acid polymorphisms (SAPs), to infer single nucleotide polymorphism (SNP) alleles. SNPs can be inherited and subsequently detected in peptides from digested protein [3]. These peptides, termed genetically variant peptides (GVPs), are especially useful in samples where DNA may be absent or highly degraded, such as is often the case with hair, fingermarks, and bone [7, 8]. Currently, mitochondrial DNA is the mainstay in HID for highly degraded samples and archaeological remains due to its multiplicity in each cell, rapid evolution, and familial information [9, 10]. Proteomic genotyping offers intrinsic advantages compared to other DNA-based genetic analyses. Protein, much like mitochondrial DNA, has many copies per cell. Processing of protein does not involve an amplification step or preliminary knowledge of the sequence, as is the case for DNA primer design. The peptide bond is chemically stable and common chemical modifications are predictable and accommodated by spectral matching algorithms [11, 12]. Protein has proven to outlast DNA due to degradation effects for archaeological sex estimation from teeth [6].

Tryptic peptides average only 14 residues, which reduces the probability of random cleavage and information loss [13]. The stoichiometry of protein copy number is greater than DNA by up to 7 orders of magnitude with a median increase of 5 orders of magnitude, allowing for detection and analysis without amplification [14–16].

Initial development of proteomic genotyping has focused on hair shafts as a source of protein-based genetic information. Over 400 genetically variant peptides from hair have been identified that can predict the corresponding SNP allele [17]. Optimization to this point has focused on chemical processing to maximize peptide production from a single hair shaft [17–19]. Aside from a shift to more modern mass spectrometry instruments for data dependent acquisition, relatively little has been done to optimize data acquisition at the mass spectrometry level. Another side effect of the current focus on data dependent acquisition for proteomic genotyping is the dependence on peptide spectral matching for detection of genetically variant peptides. Peptide spectral matches, depending on the algorithm, come with a statistical expectation score or other measures of confidence [11, 12, 20–22]. For most proteomic applications this is not functional, since multiple peptides are often identified for each gene product and the level of uncertainty can be miniscule [22]. Proteomic genotyping, however, relies on single peptides to infer SNP alleles. Validation of the inferred SNPs are therefore necessary and most easily provided by direct confirmation of genotype by DNA sequencing. Validation is also possible to confirm peptide identification through the addition of stable isotope labelled (SIL) synthetic peptides into a sample digest. These standards, equivalent in sequence and chemistry to the matching endogenous peptides, behave identically to matching endogenous peptides, and do not interfere with endogenous peptide detection. The use of SIL peptides is a standard feature of targeted mass spectrometry platforms for use in triggering endogenous peptide detection and quantification [23].

For proteomic genotyping to be readily available to forensic investigators, it also needs to be conducted on platforms that are widely accessible to investigators. Targeted mass spectrometry using triple quadrupole systems is commonly used for many forensic toxicology analyses. It is also affordable, robust, and reproducible for both chromatography and mass spectrometry. Use of targeted methods of mass spectrometry potentially improves sensitivity and bolsters analyte identification confidence, helping to fulfill the guidelines in forensic analyte identification. Guidelines set by the scientific working group for forensic toxicology (SWGTOX) [24], European Commission (directive 96/23/EC) [25], and World Anti-Doping Agency (WADA) [26] all list a minimum number of identification points to confirm the presence of a drug or other analyte. These identification points are derived from retention time windows, peak shape, and transitions and may not be satisfied using standard proteomic discovery techniques such as data dependent acquisition (DDA) alone without DNA-based verification. Currently, proteomic genotyping in forensic science has focused on the optimization of peptide production in sample preparation, and expansion to other forensically relevant tissue sources [17, 19]. These studies have relied on shotgun mass spectrometry and nano liquid chromatography coupled with orbitrap mass spectrometry and are yet to exploit useful alternative instrumental strategies available [17, 27, 28].

Here, we propose spiking SIL GVPs into hair protein digests as a means of peptide identification validation and as a mass trigger for data acquisition. This technique has typically been used for the quantification of other peptide targets [29]. A standard will elute chromatographically and be analyzed in the same time window as its corresponding endogenous GVP. Therefore, the standard and expected endogenous peptides can be directly compared in terms of retention time and ion ratios (Figures S1 and S2) providing a means for real time validation.

This study explores the capability and utility of alternative mass spectrometry platforms and data acquisition strategies using a panel of GVPs. Instead of a direct comparison of the platforms and acquisition methods which would involve an in-depth evaluation of instrument components, a proof of concept and evaluation of performance was studied for proteomic genotyping purposes only. Two approaches are first assessed: shotgun proteomics (DDA) on a Q Exactive™ Plus Hybrid Quadrupole-Orbitrap™ Mass Spectrometer, and data independent acquisition (DIA) on a Orbitrap Fusion™ Lumos™ Tribrid™ Mass Spectrometer [30, 31]. Two other approaches were also tested in the presence of matching synthetic SIL peptide standards. These include a variant of parallel reaction monitoring (PRM), called QuanDirect™ (QD) [32] conducted on the tribrid system that uses detection of SIL standard peptides to trigger data acquisition in the mass window of the corresponding endogenous peptide, and multiple-reaction monitoring (MRM) conducted on an coupled Agilent 1290/6495 triple quadrupole system [33–37]. To provide a direct comparison in the performance of genetically variant peptide (GVP) detection between the four acquisition methods, hair from European-American (n = 3) and African-American (n = 2) subjects was tested in three biological replicates. Results reported here are limited by the procedures as performed using standard mass spectrometry platforms and protocols within the range of what is considered best practice for each method. The noted enhancements, based on a panel of 24 standard peptides, have the potential to dramatically increase both the discriminatory power of proteomic genotyping and the applicability of the method since it uses instruments and analyte detection criteria commonly found in toxicology laboratories.

## 2. Experimental Section

### 2.1 General Experimental Design

Hair shafts were collected from 5 individuals who are representative of two populous ancestral backgrounds in of the United States: European and African. The number of individuals needed for this study was minimal since this was a novel proof of concept study to demonstrate the usage of targeted proteomics for proteomic genotyping. Enough donors were used to assess reproducibility and calculate standard deviation. Three single hairs from each individual, 2 cm in length, were processed separately using a previously developed method, with a total of 15 hair digests. A blank with trypsin and without trypsin were also processed in parallel with all other digests. 24 stable isotope labeled (SIL) standard genetically variant peptides (GVPs) were spiked into the hair digests for parallel reaction monitoring and multiple reaction monitoring (Table S1). Raw mass spectral data were processed using the Skyline software for the 24 GVPs of interest and their corresponding heavy-isotope peptide standards. 24 peptides were chosen to adequately represent the diversity of the full set of 408 currently identified GVPs in terms of detection sensitivity, length, and composition. Basic statistical analyses were conducted such as standard deviation, random match probability, false discovery rate (FP/(FP+TP)), and detection sensitivity (TP/(TP+FN)) to compare the three analytical methodologies. Random match probability calculations were estimated using the procedure outlined in Parker et al [3].

### 2.2 Hair Collection and Processing

Samples used in this study were prepared as part of an earlier study [17]. Briefly, five individuals were analyzed: three subjects of European (Davis, CA) and two subjects of African genetic background, respectively (Sorenson Forensics LLC, Salt Lake City, UT). Hair and saliva were collected using protocols compliant with the Institutional Review Board at the University of California, Davis (IRB# 832726). Hairs were collected by cutting a few inches inward from the distal end, therefore excluding the roots. The length of hair on the head before cutting was roughly 10 cm. Hair shafts were further cut to a length of 20 mm before continuing with protein extraction [17]. The African hair samples weighed almost half of the weight of the European hair samples due to differences in hair shaft width and shape (data not shown). Hair shafts were biochemically processed using an optimized processing protocol as part of an earlier study [17]. After initial preparation and use of the samples to generate the data-dependent acquisition datasets used in the cited study, the remaining supernatants were stored at −20°C. Prior to mass spectrometric analysis, the samples were again centrifuged to minimize insoluble particulates in the supernatant.

### 2.3 Selection of a Panel of Stable Isotope Labeled Genetically Variant Peptides

A panel of 24 highly characterized GVPs from 12 loci was selected to represent a wide range of potential sensitivities, from rarely to frequently detected, and were well characterized over a range of studies and laboratory groups [3, 17, 19, 27, 38]. The peptides selected range from 8 to 21 amino acids in length and were all modified with stable isotopes at the C-terminal lysine (+8 Da) or arginine (+10 Da) (JPT peptide technologies, Acton, MA) (Table 1). The peptides (14 nmol/well) were subsequently suspended in 4 μL of 70% formic acid and 136 μL of 0.1% trifluoroacetic acid. 10 μL of each standard were pooled to make a concentration of 4.16 pmol/μL per standard. The pooled sample was then purified using a silica C18 macrospin column with loading capacity of 30-300 μg of peptide material (The Nest Group, Southborough, MA). Briefly, the peptide digests were loaded onto the column and spun, the column was washed with 0.1% trifluoroacetic acid and spun three times, and the peptides were eluted using 80% acetonitrile 0.5% formic acid and spinning and were subsequently dried down. This pool was then injected into four hair digests from two individuals at 1 and 2 fmol/μL. A second pool was then created, with normalization based on peak area to make a final spike mixture (Table S1). This final spike solution consisted of a total of 433 nmol of peptides in 1 mL final volume (433 pmol/uL). The final mixture (1 fmol) was included in each sample injection for QuanDirect analysis and 73 pmol of the final mixture was injected into each sample for MRM analysis on the triple quadrupole (QQQ) platform.

**Table 1.**
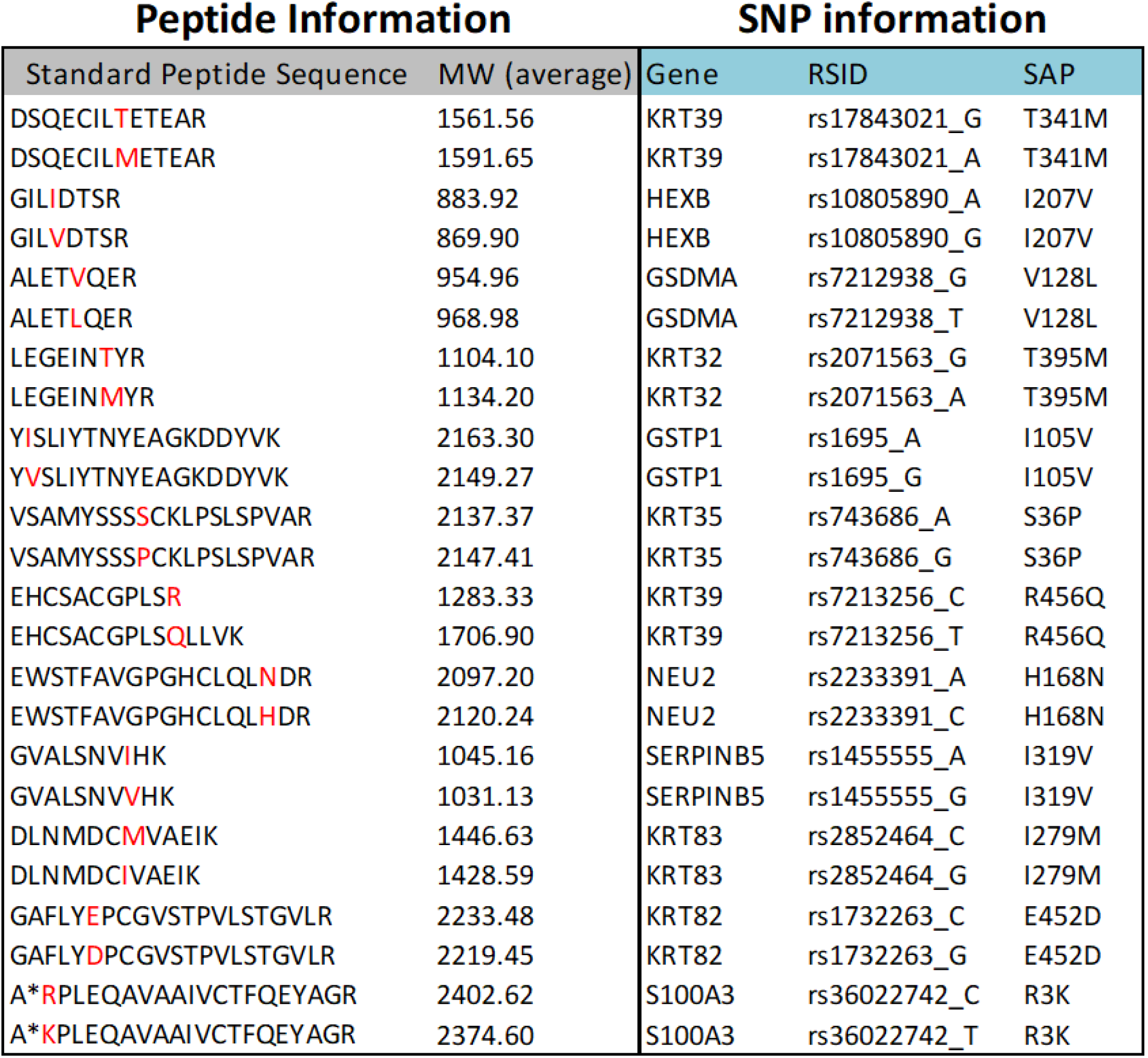
Genetically variant peptide standards. These peptides were obtained from JPT Peptide Technologies and were pooled and spiked into 17 matrices from five subjects. These were spiked only into fractions being analyzed via PRM and MRM. Red amino acids indicate the SAP location per GVP, and * indicates acetylation. All cysteines (C) are carbamidomethylated (+57), and all C-terminal amino acids are isotopically labeled. R contains 6 x ^13^C and 4 x ^15^N (+10 Da) and K contains 6 x ^13^C and 2 x ^15^N (+8 Da).

### 2.4 Instrumental Analysis

Before applying the samples to LC-MS/MS, solubilized tryptic peptides were quantified using the Pierce™ Quantitative Fluorometric Peptide Assay (ThermoFisher) as reported in previous work [17]. Resulting data were used to determine how much material to apply to the instrument. For the Q Exactive Plus and Fusion Lumos, this amounted to 0.75 μg of peptide digest material. Digest injection volume was held constant on the QQQ platform, but volumes varied based on concentration for the other three platforms.

Three instruments were used to conduct four data acquisition methods (Figure 1). The first analysis method (QE+) was conducted on a Q Exactive Plus nLC-MS/MS platform which employed data dependent acquisition (DDA). This method was established as part of an earlier study [17]. For the second analysis (Lumos-DIA), samples were analyzed on a Thermo Scientific Fusion Lumos mass spectrometer and was connected to a Dionex nano Ultimate 3000 (Thermo Scientific) with a Thermo Easy-Spray source. The acquisition method was set to data independent acquisition (DIA). For this method, peptides were trapped and separated on a 100 μm x 250 mm C18 column with 3 μm particle size PepMap Easy-Spray (Thermo Scientific) using a Dionex Ultimate 3000 nUPLC at 200nl/min. Peptides were eluted using a 90 min gradient of 0.1% formic acid (A) and 80% acetonitrile (B). Gradient conditions include 2% B to 50% B over 60 minutes, followed by a 50%-99% B in 6 minutes and then held for 3 minutes, then 99% B to 2% B in 2 minutes. The mass spectrometer was run in DIA mode using a collision energy of 35, resolution of 30K, maximum inject time of 54 ms and an automatic gain control (AGC) target of 50,000. Each individual sample was run in DIA mode with staggered isolation windows of 12 Da in the range 400-1000 m/z. For each analytic sample, the individual sample was run in DIA mode using the same settings as the chromatogram library runs except using staggered isolation windows of 8 Da in the m/z range 400-1000 m/z.

**Figure 1.**
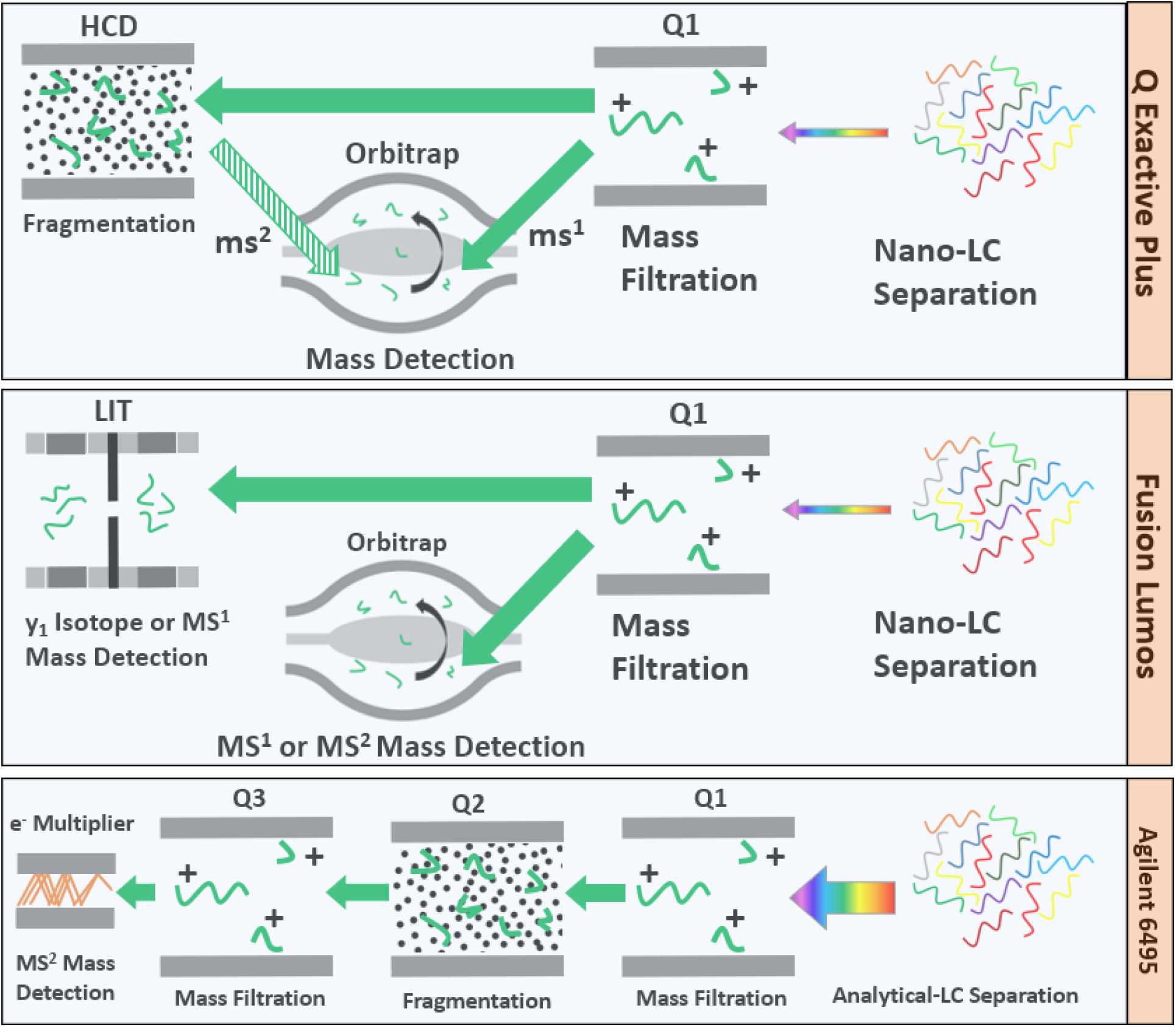
A summary of mass spectrometry data acquisition methods. 2 cm of scalp hair was digested. This peptide mixture was then analyzed using four different LC-MS/MS data acquisition methods; two methods of untargeted mass spectrometry (data-dependent acquisition on the Q Exactive plus, QE+, data independent acquisition on the Fusion Lumos, Lumos-DIA) and two methods of targeted mass spectrometry (parallel reaction monitoring on the Fusion Lumos, Lumos-QD, multiple reaction monitoring on a triple quadrupole Agilent 6495, QQQ). For the targeted methods, an isotope-labeled peptide mix was spiked into the hair digest. Both the isotope labeled peptide and endogenous peptide elute together and their MS^2^ spectra were compared to confirm the presence of the light isotope endogenous peptide.

The third analysis method (Lumos-QD) was conducted on the same instrument as the Lumos-DIA method, except QuanDirect™ (QD) parallel reaction monitoring acquisition was employed. For this method, the digested peptides were reconstituted in 2% acetonitrile/0.1% trifluoroacetic acid and 1 μg in 5 µl of each sample was loaded onto a 75 μm x 20 mm PepMap 100Å 3U trap (Thermo Fisher Scientific) where they were desalted online before being separated on a 50 μm x 150 mm 100 Å 2U PepMap EasySpray column (Thermo Fisher Scientific). Peptides were eluted using a 90 min gradient of 0.1% formic acid (A) and 80% acetonitrile (B) with a flow rate of 200nL/min. Gradient conditions include 2% to 50% B over 66 min, 50% to 99% B over 2 min, and then held at 99% B for 4 min, followed by 99% to 2% B over 2 min. Targeted precursors were interrogated for a maximum 3 sec cycle. The target list consisted of the *m/z* ratios and charge states of the heavy peptides (Table 1) only since no retention time info was required. The QuanDirect™ method of targeted mass spectrometry was used to search for endogenous GVPs [32, 39]. To trigger a data-dependent scan, the precursor must match the expected charge state and the *m/z* within 10 ppm. These precursors were interrogated with a short SRM scan (MS^2^ IT HCD) of the predicted y_1_ fragment ion region for the expected heavy R (180-190 *m/z*) or heavy K (150-160 *m/z*) peptides. The y_1_ fragment in proteomics determines the C-terminus ending and is more easily identifiable in a mass spectrum due to its monomeric independence. The ultra-fast SRM scans were performed using a 10 *m/z* mass range, rapid ion trap scan rate, HCD NCE 40%, 0.7 *m/z* isolation window, AGC target 1E4, and a max IT of 10 ms. The y_1_ ion in the SRM scan must be above an intensity threshold of 1000 and within 1 Da of the expected *m/z*. If the expected heavy y_1_ fragment ion was detected in the SRM scan, 185.1 (heavy R) or 155.1 (heavy K), then full HRAM HCD MS/MS scans were triggered on the spiked-in heavy peptide as well as the endogenous form (Table S2). These scans use the following parameters: scan range 150-1500 *m/z*, 60K resolution, 30% NCE HCD, AGC target 2e5, and max IT of 110 ms. To trigger on the endogenous peptide, an *m/z* offset of −5 or −4 U was used for the R and K peptides, respectively.

For the fourth analysis (QQQ), samples were analyzed via multiple reaction monitoring acquisition using an Agilent 1290 Infinity series HPLC system, coupled to an Agilent 6495 triple quadrupole mass spectrometer with an Agilent Jet Stream source. 10 μL of a 1:10 spike:digest (v/v) ratio (~15 μg digested material and 291 pg of spike) was loaded on a 2.1 mm × 100 mm, 2.7 μm AdvanceBio Peptide Map fused-core silica column (Agilent), and separated over a 15 min gradient at 400 μL/min.

The solvent gradient for the elution of peptides began with 5% ACN and increased to 35% ACN at 11 min, 65% ACN at 12.5 min, and 90% ACN at 13 min and held for 2 min, and then reduced to 5% for 5 min to re-equilibrate the column. Source conditions included a gas temperature at 150°C at a flow rate of 11 l/min, nebulizer pressure of 30 psi, sheath gas temperature of 150°C at 10 l/min, and a capillary voltage of 3500 V. Collision energies were calculated based on precursor m/z and charge state in Skyline software, and were not fully optimized. Data were acquired in positive dynamic MRM mode (dMRM) with an MS^1^ resolution set to wide and MS^2^ resolution to unit, retention time window of 30 sec, and a cycle time of 500 ms. Three transitions were selected for the detection of each standard and endogenous peptide (Table S3).

### 2.5 Software Analysis

To make the resulting QuanDirect™ PRM datafiles amenable with Skyline software, spectra from the linear ion trap were excised using the FT RecalOffline tool from Xcalibur™ (ThermoFisher Inc.). This treatment does not interfere with spectrum interpretation since this was only a part of the internal decision tree. Raw files were manually loaded and the external slicer was called, under Rawfile Functions, from within RecalOffline to remove any masses below 200 *m/z* using a mass filter from 200 *m/z* to 2000 *m/z*. Since only low mass y_1_ fragments of 185 and 155 *m/z* with scan range < 200 *m/z* were searched for, the filter removed all of the ion trap data from the file. The resulting file only contained MS^1^ and MS^2^ orbitrap data which was used to analyze endogenous and standard peptides.

Skyline software [40] (version 20.1) was used to visualize endogenous peptide data and SIL peptide data simultaneously and to extract only mass transitions of interest (Tables S2 and S3). Positive peptide identification required a s/n ratio > 3, peak intensity of ion targets > 20 counts, and an ion ratio between quantifier and qualifier ions within 25% of the target. These parameters were chosen to meet minimum identification points from common forensic guidelines [24–26]. For the analysis performed here, samples were analyzed in batches based on the instrumentation on which they were run. For samples analyzed on the QE+ using DDA, full-scan transition settings for MS^1^ filtering were set to include count isotope peaks, orbitrap precursor mass analyzer, with 3 peaks and a resolving power set to 60,000 at 400 *m/z*. MS/MS filtering settings were set to DDA as the acquisition method, orbitrap product mass analyzer, no isolation scheme, a resolving power set to 60,000 at 400 *m/z*. All other settings were set to default. For samples analyzed using Lumos-DIA, the same settings were used as DDA with the exception of 70,000 MS^1^ resolving power, DIA as the acquisition method, results only as the isolation scheme, and 17,500 as the MS/MS resolving power. For samples analyzed using Lumos-QD, the same settings were used as Lumos-DIA except for no isotope peaks, MS/MS filtering settings were set to targeted as the acquisition method, and no isolation scheme. For samples analyzed using QQQ with MRM, the same settings were used as PRM except for MS^1^ filtering were set to include count isotope peaks. One precursor and 3 transitions were chosen for MRM analysis, while one precursor and 10-24 transitions were chosen for the PRM analysis and DDA analysis. These are both above the minimum standard guideline for the number of ions required for a positive identification [25, 26].

Positive peptide identifications were called from the Skyline software based on precursor and transition signal to noise ratio, retention time, transition masses, and ion ratios. For retention time, this identification criteria included having a GVP retention time within 2% or ± 0.1 min of the labeled standard. For the DIA and DDA approaches, comparison of retention time to a labeled standard was not used. In terms of signal to noise ratio, a minimum ratio of 3:1 was used as the threshold for data from all platforms. Transitions used for identification for the targeted approaches were taken from the most abundant transitions for the labeled standard peptides. The untargeted methods utilized the Prosit library [41] to compare both transitions for identification and ion ratios. For all acquisition methodologies, ion ratio maximum tolerance windows were set to be within 10% of the relative abundance of the compared ion, so long as the peak is at least 50% of the base peak [26]. The PRM and MRM methods used the labeled standard peptides as a reference and the DDA and DIA methods used the Prosit library as the reference for ion ratios.

### 2.6 Statistical Analysis

GVP Finder (v1.2) (https://www.parkerlab.ucdavis.edu) was used to estimate random match probabilities (RMPs). This is an excel spreadsheet compatible with X!Tandem output developed in previous work [17]. In short, RMP was calculated using the product rule [3, 42] by simply multiplying independent genotypic frequencies based on observations on individual genotypes from the major populations in the 1000 Genome Project Consortium [43]. To account for linkage disequilibrium, it was assumed that there was complete linkage for GVPs shared within an open reading frame and complete independence between each open reading frame [3]. For GVPs that were determined to be genetically linked within an open reading frame, a cumulative genotypic frequency was calculated by counting the number of individuals in the consortium who have the same gene specific GVP profile as was obtained from the sample and dividing by the total number of individuals in the population. Genetic validation was performed to assign trueness of positive and negative detections. Genomic DNA was extracted and sequenced as reported in previous work [17].

To estimate random match probability of a profile that would result from a targeted QQQ analysis using all known GVPs, and assuming equivalent detection sensitivity obtained from the panel, a Markov Chain Monte Carlo (MCMC) model was developed. MCMC is an algorithm that simulates stochastic processes such as sampling from a probability distribution [44, 45]. This method of sampling allows an estimation of true population probability distributions by randomly sampling from probabilistic data. For this study, MCMC was developed as a function of GVP number and validated by superimposing actual RMP values from previous studies. The probability distributions were taken from actual genotype frequencies from the 408 GVPs that have been identified. A theoretical genotype of non-synonymous SNP alleles was generated based on randomly selecting known GVPs and randomly determining if a theoretical genotype would include that GVP based off its genotype frequency. One hundred iterations were included in this model. Minimum, maximum, and median theoretical RMP values were estimated based on which theoretical GVPs were randomly chosen in the model. The model assumes one GVP locus per open reading frame. The resulting modelled genotypes were randomly selected in each iteration as a function of prior probability based on the genotype frequency chosen randomly here, rather than favoring GVPs of historically higher detection. Therefore, this model is not biased towards specific GVPs and does not mimic biological GVP profile distributions.

### 2.7 Data Reporting and Availability

All RAW data files containing detected endogenous peptides and SIL GVPs from hair digests mentioned in this work, including from the supplemental section, are publicly available on ProteomeXchange (PXD024651) [46]. A complete list of datafiles is also available (Table S4). Files from QE+ are comprehensive and include all detected ions, whereas the Lumos-QD and QQQ files are limited to ions corresponding to GVPs from the panel. Lumos-QD data are modified to exclude linear ion trap data. Skyline files for data obtained from the four analytical platforms are publicly available at https://panoramaweb.org/TargetedGVP.url [47].

## 3. Results and Discussion

Studies optimizing the detection of genetically variant peptides (GVPs) have done so by focusing on the chemical release of tryptic peptides from the hair matrix, or by applying the resulting peptide mixtures to more sensitive instrumentation. In this study different mass spectrometry data acquisition methods were tested to evaluate additional options for increased GVP detection and therefore further increase the utility of proteomic genotyping in forensic investigation. Accordingly, mass spectrometry data acquisition using Data Independent Acquisition (DIA), Parallel Reaction Monitoring (PRM) and Multiple Reaction Monitoring (MRM) were all tested on instruments with configurations that are standard for each method. Acquired data from all three platforms, as well as existing data using a shotgun proteomics Data Dependent Acquisition (DDA) approach, were screened for detection of a panel of 24 endogenous GVPs in replicate trypsin digests using a common bioinformatic workflow in Skyline (Figures 2, 3, and S1). The cumulative inferred non-synonymous SNP genotypes were directly validated using the exome of each subject (Figure 3) to determine basic metrics such as true positive, false positive, true negative, and false negative rates, along with sensitivity (TP/(TP+FN)) and false discovery rates (FP/(FP+TP)). Besides this binary classification process, other metrics such as signal to noise, peak shape, ion ratio, peptide ionization efficiency, retention time, and abundant transitions were also measured. In the case of the PRM and MRM acquisition methods, the evaluation was facilitated by using a panel of exogenous stable isotope labeled peptides. While each data acquisition approach was within the range of normal best practice, no systematic optimization occurred beyond establishing basic acquisition and chromatographic parameters. The results therefore reflect different chromatographic, ionization, and mass spectrometer systems and configurations for each acquisition method. While direct comparisons could not be made, the performance of each method could be individually evaluated in comparison to previously acquired data using shotgun proteomics (DDA).

**Figure 2.**
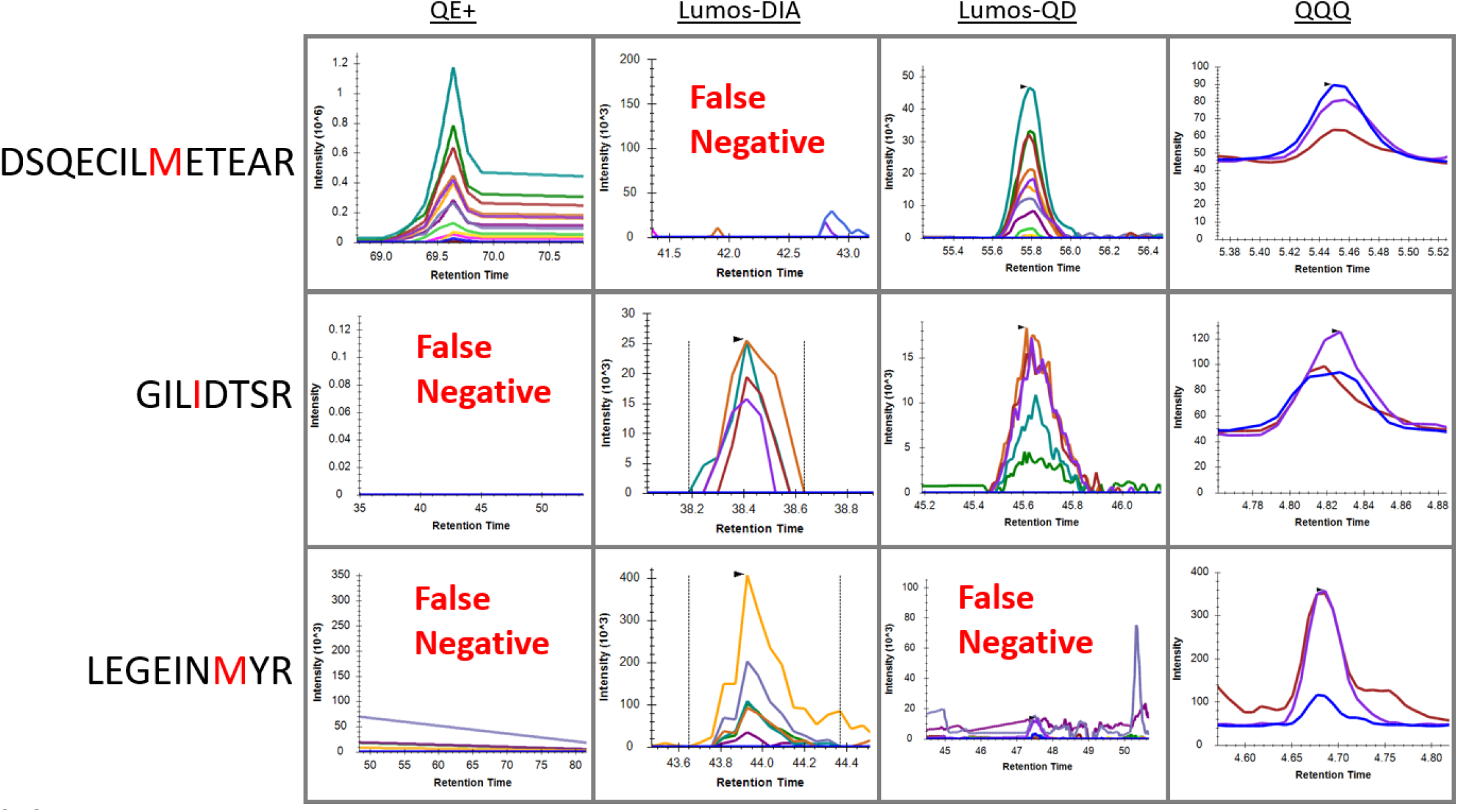
Performance of four analytical platforms. This figure demonstrates the usefulness of targeted proteomic methods for three of the 24 GVP peptides analyzed. All peptides are expected to be present in the sample as confirmed by genotyping. However, the first peptide (DSQECILMETEAR) is missing in the Lumos-DIA, the second peptide (GILIDTSR) is missing in the QE+, and the third peptide (LEGEINMYR) is missing in both QE+ and Lumos-QD.

**Figure 3.**
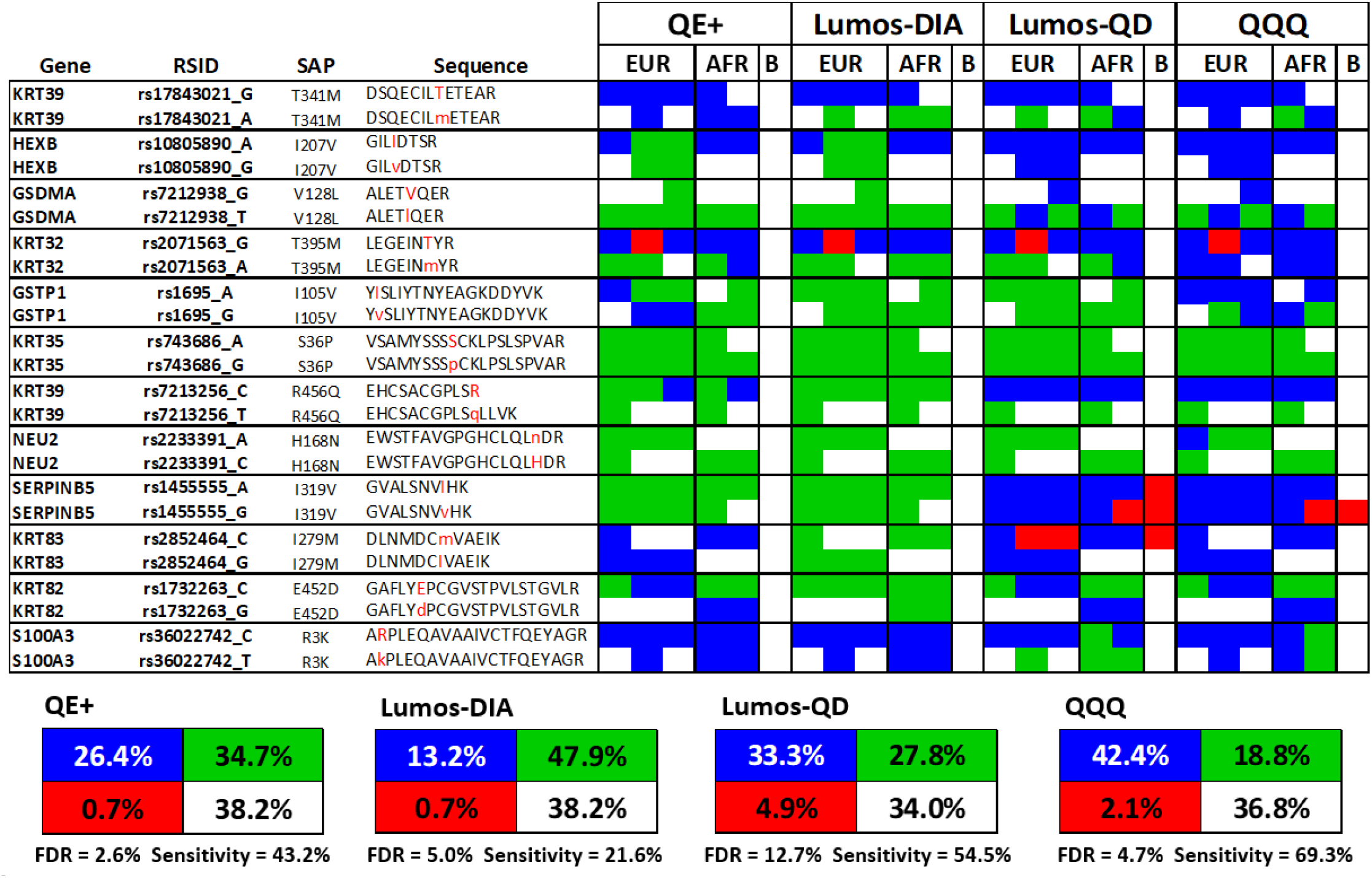
GVP matrix evaluating four analytical methods. This matrix represents GVPs that have been verified via whole exome sequencing. Each row is a variant peptide and each column is an accumulated GVP profile from three replicates. QE+, data dependent acquisition on Q Exactive+; Lumos-DIA, data independent acquisition on the Fusion Lumos; Lumos-QD, QuanDirect on Fusion Lumos; QQQ, multiple reaction monitoring on Agilent 6495; EUR, three European subjects; AFR, two African subjects; TP, true positive; FN, false negative; FP, false positive; TN, true negative; FDR, false discovery rate.

### 3.1 Analysis of Performance for the QE+ Platform

Data previously acquired on a nano-LC / Q Exactive Plus (QE+) platform with data dependent acquisition (DDA) was reanalyzed using Skyline software (version 20.1) to provide a benchmark for other data acquisition strategies and instrument configurations [17, 40]. The percentage of true positive shotgun proteomic identifications for the QE+ platform was 26.4% and the detection sensitivity (TP/(TP+FN)) was 43.2% (Figure 3). The false discovery rate (FP/(FP+TP)) was the lowest for the QE+ platform at 2.6% and the average GVP detection from the 24-GVP panel was 6.5 ± 1.1 (Figure 3). Ten of the 24 endogenous peptides were not detected at all using this platform. These peptides include GILVDTSR, ALETVQER, ALETLQER, EWSTFAVGPGHCLQLNDR, GVALSNVIHK, and GVALSNVVHK from proteins HEXB, GSDMA, NEU2, and SERPINB5 (Figures 3 and S3). These peptides may not have been observed due to low protein abundance in the hair sample digests, whereas keratin proteins are very abundant [48]. No peptides were observed using this method that were not also observed in the other methods. Four endogenous peptides were not detected at all in this study regardless of the platform used: VSAMYSSSSCKLPSLSPVAR, VSAMYSSSPCKLPSLSPVAR, EHCSACGPLSQLLVK, and EWSTFAVGPGHCLQLHDR. These longer peptides may be more challenging to detect based on their length and residue composition [49]. For a positive detection, the average signal to noise ratio was higher than 1:3 (Figure 2, S5), defined as the variance of amplitude of the baseline and signal was the amplitude of the peak as measured from the apex to the average baseline, which meets the minimum requirements in toxicology scientific working group guidelines [24–26]. The nanoflow chromatography / Q-Exactive configuration used in this analysis was sensitive [50], requiring only 1 µg of the roughly 100 μg protein present in 2 cm of a single hair shaft [51]. This acquisition method can be used for GVP discovery and provide a resource for retrospective analysis of GVPs, although the analysis was limited to the 24 GVPs and associated ions in the SIL-peptide panel. In terms of the peak shape for the QE+ analytical platform, the overall form differs from the three other platforms due to differences in chromatography (Figure 2). Complex chromatography patterns due to internal prolines were detected [52, 53].

### 3.2 Analysis of Performance for the Lumos-DIA Platform

For Data Independent Acquisition (DIA), samples were applied to a Dionex nano Ultimate 3000 coupled to an Orbitrap Fusion™ Lumos™ Tribrid™ Mass Spectrometer and data acquired using SWATH-MS (DIA) (Figure 1). Overall, Lumos-DIA peaks appear sharp with peak intensities averaging at 1 × 10^3^ and the average signal to noise ratio was also above 1:10 (Figure 2). The resulting percentage of true positive identifications for Lumos-DIA was 13.2% and the false discovery rate was 5.0% (Figure 3).

Average GVP detection from the 24-GVP panel was rather low, at 3.1 ± 0.8 and therefore detection sensitivity (TP/(TP+FN)) was also low, at 21.6% (Figure 3 and S3) which is less than half the sensitivity we typically achieve using standard DDA methodologies on the QE+. Overall, this method did not detect 15 of the 20 peptides that were detected using the other methodologies. Seven of these 15 missing peptides were found in all three other methodologies, which include peptides DSQECILmETEAR, LEGEINmRY, EHCSACGPLSR, DLNMDCmVAEIK, DLNMDCIVAEIK, GAFLYEPCGVSTPVLSTGVLR, and GAFLYdPCGVSTPVLSTGVLR from KRT39, KRT32, KRT39, KRT83, and KRT82 (Figures 3 and S3). Only one peptide was observed using the Lumos-DIA methods that was not observed in either PRM or MRM method for the same donor; AKPLEQAVAAIVCTFQEYAGR.

DIA requires little method optimization [30, 54], and may be used for GVP scouting or retrospective use. Unbiased detection of peptides, uniquely for DIA, allows for proteins of low abundance to be detected [31]. This precludes the need for an exclusion list or other mass filtering parameter optimizations. The data is highly reproduceable [31] and so running evidence samples alongside exemplars would be more consistent and would result in less variance due to protein abundance levels [55]. The main challenge in DIA interpretation, at least in our hands, was deconvolution of MS^2^ spectra. In this data false positive identification occurred in four peptides that were not explained by instrument carry-over or genetics. Due to the nature of SWATH mass spectrometry, an MS^2^ mass spectrum may contain product ions from multiple precursor ions, which may lead to convoluted MS^2^ extracted ion chromatograms. This may be problematic in a courtroom setting, although the use of the internal standard SIL peptides would have significantly aided MS2 interpretation. The sensitivity of this analysis may also be improved with better precursor validation, library match validation, and staggered SWATH windows [56].

### 3.3 Analysis of Performance for the Lumos-QD Platform

Targeted parallel reaction monitoring (PRM) was evaluated by analysis of replicate digests using an analytical variant called QuanDirect™ (QD) (Figure 1) [32]. This method differs from classical PRM by triggering data acquisition using detection of the y_1_ SIL amino acid instead of characterized retention times. The resulting percentage of true positive identifications for Lumos-QD was 33.3% and the false discovery rate was also highest, at 12.7% (Figure 3). The high false discovery rate was due to higher levels of carry-over of the peptides GVALSNVIHK, GVALSNVVHK, and DLNMDCMVAEIK, including in the blanks. Average GVP detection from the 24-GVP panel was 9.5 ± 0.7 and detection sensitivity (TP/(TP+FN)) was 54.5% (Figure 3 and S3). Overall, the methodologies we applied to the Lumos-QD platform revealed a 1.3-fold increase in detection sensitivity in comparison to the traditional methodologies we applied to the QE+ platform (Figure 3). However, the peptides ARPLEQAVAAIVCTFQEYAGR and AKPLEQAVAAIVCTFQEYAGR were both detected inconsistently in the Lumos-QD series when compared to the three other methods.

Overall, Lumos-QD peaks appear sharp and symmetrical with peak intensities that averaged at three orders of magnitude and an ion current signal to noise ratio above 1:10 (Figure 2). As expected, peaks identified using Lumos-QD were less stable in retention time. Standard peptide peaks drifted between runs by an average variance of 20 sec (or 0.4% of total run time), which is longer than the average peak width of 15 sec (or 0.3% total run time) (Figure S2). Peak variance was on average 1.5x larger for the Lumos-QD method compared to the QQQ method, described below. Peak drift between a standard peptide and its corresponding endogenous peptide in each run was minimal for both PRM and MRM analyses (Figure S2B/C).

The QuanDirect method addresses a major weakness of PRM, namely retention time variability that results from low flow chromatography. The more recent SureQuant methodology [57] continues this mass triggering approach by searching for the internal standard precursor ion in a fast and low-resolution watch mode and switches to a high-resolution quantitative mode when the isotope-labeled precursor ion is detected [57]. For QD, not having to pre-determine strict elution windows saves time and effort. However, the traditional PRM methodology when it is optimized, and takes full advantages of SIL characteristics and instrument cycle windows, and may be more sensitive and optimized. The PRM acquisition method is easier to establish since it is not limited by prior identification of targeted transitions. Of course, targeted acquisition only acquires limited information and therefore cannot be used retrospectively.

### 3.4 Analysis of Performance for the QQQ Platform

Multiple reaction monitoring (MRM) was conducted using an Agilent 1290 Infinity series HPLC system coupled to an Agilent 6495 triple quadrupole mass spectrometer (Figure 1). In this targeted acquisition experiment, a panel of 24 SIL GVPs (Table 1, Figure S4) was added to provide a direct comparison of transition signals and retention times for endogenous GVPs. The resulting percentage of true positive identification was 42.4% and the false discovery rate (FP/(FP+TP)) was 4.7% (Figure 3).

Average GVP detection from the 24-GVP panel was 11.7 ± 1.7 and detection sensitivity (TP/(TP+FN)) increased to 69.3% (Figure 3 and S3). Overall, the methodologies we applied to the QQQ platform revealed a 1.6-fold increase in detection sensitivity in comparison to the traditional methodology we applied to the QE+ platform. Peak shape was more uniform in the QQQ run where retention time drifted between runs by an average variance of 1.5 sec, which is shorter than the average peak width (4.8 sec) (Figure S2A). Peak drift between a standard peptide and its corresponding endogenous peptide in each run was minimal (Figure S2B/C). The average signal to noise ratio for QQQ was lower and the overall detected peak intensities were lower by an average of 3 orders of magnitude.

The QQQ method resulted in noisier peaks due to the smaller number of transitions selected, lower mass accuracy, and shorter run times that may have resulted in overlap with extraneous ions.

However, the QQQ system provided the greatest increase in sensitivity. In terms of method development, MRM on a QQQ system requires more development to deal with limited selection of transition masses, detection parameters and manual optimization of acquisition parameters such as collision energy, retention windows, dwell time, duty cycle, and cycle time. In terms of input material, the QE+, Lumos-DIA, and Lumos-QD methods are all the same, with 1 μg of material injected due to their use of nano-LC. However, the QQQ system used a volume of 10 μL, which averaged to ~15 μg of digested peptide material. This is a 15-fold increase in starting material, but still only 10 to 20% of a protein digest from a single hair shaft (20 mm). The MRM method depends on a limited number of transitions for identification. The performance of each transition therefore needs to be individually evaluated and alternative transitions selected as necessary. Selection of the target GVP for a given non-synonymous SNP is also a major consideration. The current approach to proteomic genotyping is based on shotgun proteomics that allows genotype inference to occur from several chemical variants, or ‘peptidoforms’, of a GVP [31, 58, 59]. These result from expected but variable environmental chemical modifications such as deaminidation, methionine oxidation and N-terminal acetylation. Selection of a representative peptide will ideally occur from the peptidoform with the highest signal from samples derived from a range of real-world contexts.

The false positives identified in this study have three potential causes. The first category is genetic. This class of false positive is demonstrated by the peptide in K32 protein containing the SAP T395M. An uncommon variant in K40 (W390R) results in the same genetically variant peptide sequence, which was positively identified in subject E2. As described earlier, the second class of false positive detection is due to instrument carry-over. SerpinB5 and K83 found in Lumos-QD may exemplify this since these were also found in the method blanks and not in reagent blanks. These most likely reflect instrument carry-over and not reagent contamination due to the low peptide abundance. The associated peaks are smaller than that observed in other true positive hair shaft digests by more than two orders of magnitude (data not shown). The degree of carry-over for any peptide marker can be factored into appropriate thresholds during the development process for designating a positive detection of endogenous GVP [24–26]. Assessment of inter-sample blanks is a crucial step which must be included. Caution should be used to avoid a third category of false positive detection involving data interpretation.

The presence of false positive assignments, or potential assignments, raises the issue of peptide validation and what constitutes a positive determination. In targeted proteomics, positive determination is more straightforward. If the retention time, precursor ion ratios, product ion ratios, and mass errors are consistent to those of the SIL standard within a certain range, then the peptide is positively assigned. If one of these aspects is missing, then it is still possible to validate through the other measures. For example, precursor ions are missing for many GVPs in the Lumos-QD analysis including HEXB 207I, GSDMA 128L, and K39 456R. However, other measurements such as product ion ratios (data not shown) and retention times (Figure S2) are consistent with the standard. Therefore, these are considered positive assignments. Without the use of SIL standards, as we see with the QE+ and Lumos-DIA methods, this is not a straightforward task. As a first step of validation, a library may be used to compare precursor and product ions. In this analysis, we use Prosit [41] as a guideline for ion ratio comparison.

Traditional QQQ platforms differ significantly from research mass spectrometry platforms in chromatography and ionization. The QQQ platform used here employs an analytical column with dimensions of 2.1 mm x 100 mm and a flowrate of 400 μL/min. This platform also employs an Agilent jet stream ion source, which offers improved instrumental sensitivity, but is not as sensitive as nanospray sources. This is in comparison to the nano-LC systems of QE+ and Lumos which used column dimensions of sub-100 μm diameter and 150-250 mm lengths and a flow rates of 200-300 nL/min. These smaller diameter columns with slower flow rates offer enhanced instrumental sensitivity due to entering the column in a more concentrated band, therefore lessening radial dilution [50]. Using nanoflow columns, ion suppression effects and scan rate limitations are reduced, and the system is more responsive to temperature changes. When considering time efficiency, analytical columns offer the advantage of 15-minute proteomic runs while the nanoflow systems offer around 90 minute runs. In a non-research environment where time is crucial, and especially when dealing with forensic samples, the difference of six to nine samples run in 90 minutes versus one sample in 90 minutes can make a large difference in time efficiency and dramatically reduce the instrumentation costs per sample. Instrument downtime due to maintenance and complications is also typically lower for QQQ systems.

Meeting the requirements of the forensic science community is an important challenge in this research. The Daubert standard requires forensic evidence to meet five major milestones including testing in real-world scenarios, publication and peer review, known error rates, standards to control the technique’s operation, and general acceptance within the forensic science community [60]. SWGDAM developmental validation guidelines for genetic studies are similar, with objectives including characterization of genetic markers, species specificity, sensitivity studies, stability studies, precision and accuracy, case-type samples, and population studies [61]. SWGTOX gives even further guidelines on mass spectrometry standards including assessments on bias and precision, calibration models, instrument carry-over, inference studies, ionization suppression and enhancement, limit of detection, and limit of quantitation [24]. The proposed methods in this research meet both practical and legal standards. In terms of meeting the Daubert standard, GVP analysis has also undergone testing and validation studies using real-world scenarios, such as pigmentation status, body site origin and time in storage, peer review, and reported error [3, 17, 27, 62, 63]. This research contributes further by establishing the use of peptide standards as an additional validation option to investigators [3, 17, 19, 27, 38, 64, 65]. To meet the SWGDAM developmental guidelines, GVP DNA markers have been characterized, species specificity is checked, sensitivity is currently being studied, stability of peptides has been demonstrated, precision and accuracy are reported in proteomic datasets, and population studies are currently being conducted [3, 19, 27, 38, 65]. Of the targeted mass spectrometry approaches taken, the QQQ mass selection windows for primary and transition ions are broader and less selective than those used in parallel reaction monitoring. However, any broadening of specificity is more than compensated for by the consistency of retention time, particularly in the presence of stable isotope labelled (SIL) peptides. To meet SWGTOX guidelines calibration verifications, proteomic calibration models, instrument carry-over criteria are being assessed and in development, or are in place. Likewise, the use of exogenous SIL peptides for inference of endogenous GVPs using transition ion and signal to noise ratios can be reported and available for replication (Figures S1 and S5). Additional levels of validation such as establishing limits of detection and quantification are currently under investigation.

### 3.5 Extrapolation of Random Match Probability

Random match probabilities for the 24 peptide panel do not exceed 1 in 1000, which is to be expected of a small panel. However, random match probabilities have been reported to reach up to 1 in 624 million from 77 detected GVPs from a single hair shaft [17]. To model what RMP estimates could be if more sensitive targeted methods were applied, inferred genotypes were modeled as a function of increasing detection of GVPs. The modeled genotype frequencies of each allele were randomly selected from existing GVP genotype frequencies in the European reference population of the 1000 Genomes Project Consortium for 10 to 300 possible GVP detections [43]. One hundred iterations were completed and minimum, median, and maximum 1/RMP estimates were plotted for increasing GVP levels (Figure 4). Previous data from single hair and 4mg hair digests were overlaid to validate a portion of the model [17]. The model demonstrates wide variation in potential 1/RMP values and different numbers of observed GVPs that reflect the stochastic nature of inferred genotypes from randomized alleles; not every genotype contains an allele, and genotype frequencies can vary widely. Not all of the actual overlaid 1/RMP values were within the minimum and maximum boundaries for estimated RMP, which reflects a higher number of GVPs occurring within an open reading frame, and therefore were treated as a single locus when processing actual GVP profiles [3]. The contingency of multiple GVPs in an open reading frame were not incorporated into the model. Likewise, heterozygosity was also not incorporated into the model, although the resulting product of genotype frequencies of two alleles (*gf*_AB_ = *gf*_A_ x *gf*_B_) closely approximates and is slightly more conservative than the actual genotype frequency (*gf*_AB_ = 2AB) (Figure S6). The maximum difference between the two equations was only 6.25% at an allelic frequency of 0.5 (Figure S6). Expected 1/RMP values from a projected 1.6-fold increase in detection sensitivity was indicated on the model as was observed in MRM. For an estimated 130 GVP detections, projected values would range from an estimated maximum of 1 in 10^18^ (1 in 1 quintillion), to a minimum of 1 in 10^10^ (1 in 10 billion), with a median of 1 in 10^13^ (1 in 10 trillion). For a 1.3-fold increase in sensitivity, as observed using the QuanDirect variant of PRM, the roughly 100 GVP detections. This was a significant improvement to current standards in proteomic genotyping and predicts that individualization can routinely be obtained using a single human hair shaft. Based on 20 repeated iterations there was an increase in median RMP of an order of magnitude per 8.8 ± 0.9 GVP detections.

**Figure 4.**
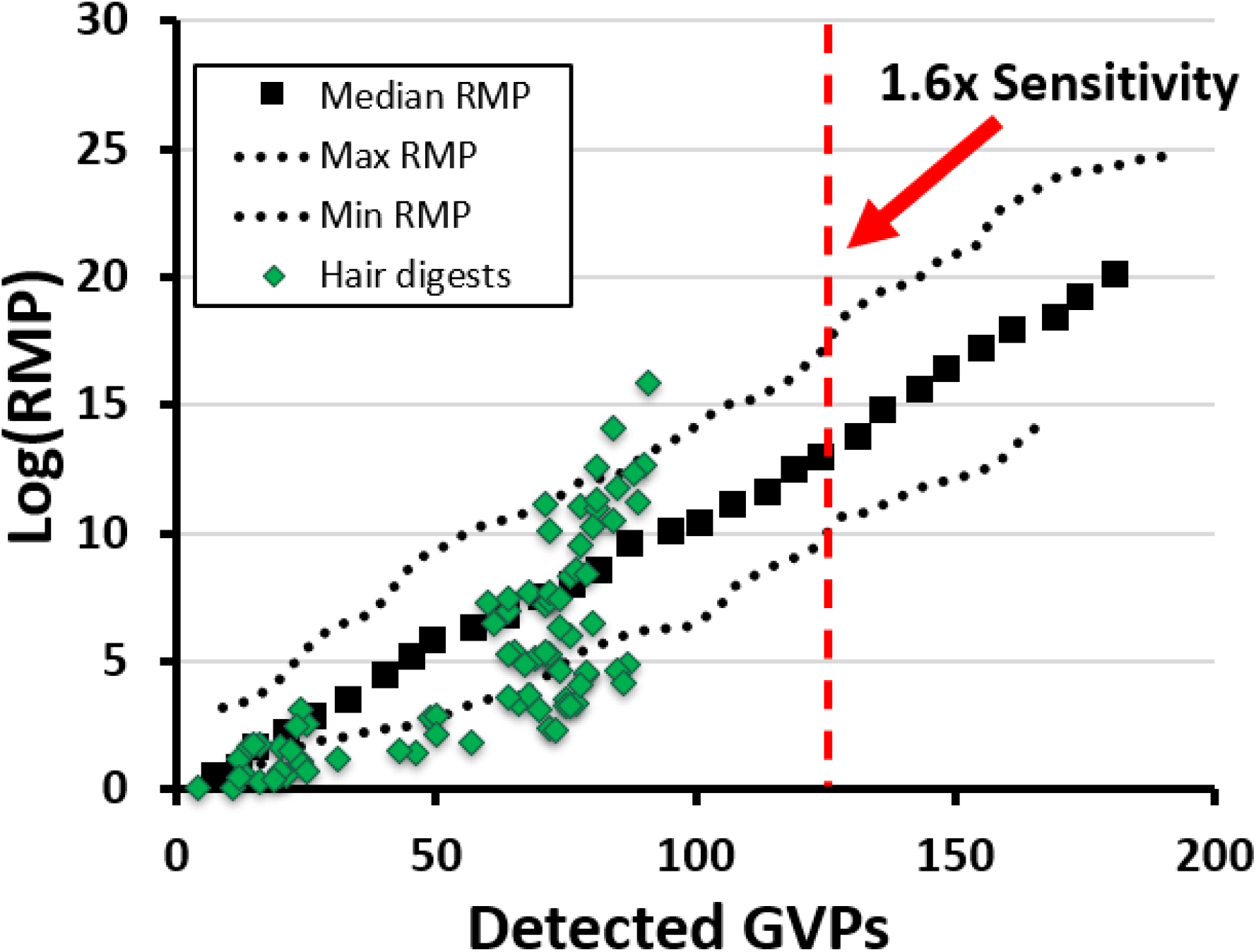
MCMC model for RMP extrapolation. A Markov Chain Monte Carlo model was developed to estimate random match probability as a function of GVP detection. After 100 iterations, the maximum, minimum, and median values were obtained. RMP values were validated from digests of 2 cm of hair shaft. QQQ detection is estimated to increase GVP detection by 1.6-fold and QuanDirect™ detection is estimated to increase GVP detection by 1.3-fold.

Many assumptions were made in this model, which elicit broad estimates of RMP. The model used does not perfectly reflect the method actually used to determine RMP from detected GVPs. GVPs from the hair shaft are often clustered in the same gene product and effects of linkage disequilibrium, accommodated in actual RMP estimates, were not taken into consideration. These differences may help to explain deviations between modelled RMPs and actual values. Actual values of RMP shown in the MCMC model are from previously published data that incorporate linkage disequilibrium into the RMP calculation [17]. The panel of 24 synthetic stable isotope labeled (SIL) GVPs used in this study were selected to represent a range of relative abundances from very frequently to rarely observed. This was to ensure that changes in detection sensitivity would be reflected in the data. They are not a random selection from the more than 400 validated GVPs identified to date and the observed 1.6-fold increase in sensitivity therefore is contingent. The increase in detection sensitivity was associated with an increase in false discovery rate, a scenario that is common in analytical chemistry. Since the panel of 24 GVPs chosen does not accurately represent the full set of 408 GVPs, a bias towards more discriminating RMPs may exist. Mitigation of this effect was attempted by choosing GVPs that vary in detection sensitivity, length, and detection. Heterozygosity also was not incorporated into the model, and since this results in a slightly more conservative RMP estimates, this may account for some of the actual 1/RMP values being more discriminating than the model.

## 4. Conclusion

This work demonstrates the utility for alternative analytical platforms in proteomic genotyping and establishes validation methods for the evaluation of inferred genotypes. Sample limitation, a lack of opportunity for reproducibility, and more stringent criteria for peptide identification are all relevant when interpreting data and communicating findings in a legal context. Maximizing relevant peptide signals is critical. Previous proteomic optimization has occurred at the level of sample processing to increase the release of detectable peptides from the hair matrix [17]. This study further optimized the detection of genetically variant peptides by focusing on the analytical framework. A range of three basic mass spectrometry approaches were utilized and associated to a reanalysis of GVP detection using standard shotgun proteomics [17]. These approaches included data dependent acquisition (DDA), systematic data independent acquisition (DIA), parallel reaction monitoring (PRM), and multiple reaction monitoring (MRM). PRM and MRM methods of acquisition also included the addition of a panel of 24 stable isotope-labeled peptides to facilitate and validate GVP detection. While each method was conducted on mass spectrometry platforms with suitable configurations for each method, additional optimizations could still be conducted for each approach, particularly DIA. Nevertheless, the MRM method performed best in terms of GVP detection, with an overall increase in detection sensitivity of 1.6x when compared to the traditional data dependent acquisition approach on a QE+ platform. This platform incorporated more robust analytical column chromatography and triple quadrupole mass spectrometry. In addition to increased sensitivity and a simplified analytical process, the ease of explanation in a legal setting, and use of preestablished methods and accreditation standards currently used in forensic toxicology should facilitate incorporation into the forensic community. In this study, targeted methods applied to GVP detection enhanced the use of hair protein as a source of human individualization, with a projected random match probability of 1 in 10 trillion if this method were applied to all 408 currently identified GVPs. Detection of human- or fluid-identifying peptides currently relies on MRM on triple quadrupole mass spectrometry platforms. An expansion of this targeted approach to include GVPs has the potential to dramatically improve the accessibility of proteomic genotyping, reducing costs and simplifying interpretation. Increased detection sensitivity will increase the discrimination and therefore utility of resulting random match probabilities. The use of targeted mass spectrometry may well place proteomic genotyping as a more accessible, quantitative, and legally explainable tool.

## Ethical Approval

All procedures performed in studies involving human participants were in accordance with the ethical standards of the institutional and/or national research committee and with the 1964 Helsinki declaration and its later amendments or comparable ethical standards. All samples were collected following the guidelines provided by the Institutional Review Board (IRB# 832726) and Institutional Biosafety Committee (IBC) of the University of California, Davis, CA.

## Acknowledgments

The authors thank Dr. Bob Rice, Dr. Blythe Durbin-Johnson, Dr. Ben Moeller, and Dr. John Newman for their advice. This publication was made possible, in part, with support from the UC Davis Genome Center Bioinformatics Core Facility. The sequencing was carried out at the DNA Technologies and Expression Analysis Cores at the UC Davis Genome Center, LC-MS/MS was supported by NIH Shared Instrumentation Grant 1S10OD010786-01. We specifically acknowledge the assistance of Jie Li, Emily Kumimoto, Siranoosh Ashtari, Vanessa K Rashbrook, and Lutz Froenicke.

## Abbreviations

DDA: data dependent acquisition
DIA: data independent acquisition
GVP: genetically variant peptide
HID: human identification
MRM: multiple reaction monitoring
PRM: parallel reaction monitoring
QD: QuanDirect
QQQ: triple quadrupole
RMP: random match probability
SAP: single amino acid polymorphism
SIL: stable isotope labeled
SNP: single nucleotide polymorphism

## References

[1] K. M. Legg, R. Powell, N. Reisdorph, R. Reisdorph, and P. B. Danielson, “Discovery of highly specific protein markers for the identification of biological stains,” Electrophoresis, vol. 35, no. 21–22, pp. 3069–3078, 2014, doi: 10.1002/elps.201400125.

[2] A. Tailor, J. C. Waddington, X. Meng, and B. K. Park, “Mass Spectrometric and Functional Aspects of Drug-Protein Conjugation,” Chem. Res. Toxicol., vol. 29, no. 12, pp. 1912– 1935, 2016, doi: 10.1021/acs.chemrestox.6b00147.

[3] G. J. Parker et al., “Demonstration of protein-based human identification using the hair shaft proteome,” PLoS One, vol. 11, no. 9, pp. 1–26, 2016, doi: 10.1371/journal.pone.0160653.

[4] H. Yang, B. Zhou, M. Prinz, and D. Siegel, “Proteomic analysis of menstrual blood,” Mol. Cell. Proteomics, vol. 11, no. 10, pp. 1024–1035, 2012, doi: 10.1074/mcp.M112.018390.

[5] K. M. Legg, L. M. Labay, S. S. Aiken, and B. K. Logan, “Validation of a Fully Automated Immunoaffinity Workflow for the Detection and Quantification of Insulin Analogs by LC-MS-MS in Postmortem Vitreous Humor,” J. Anal. Toxicol., vol. 43, no. 7, pp. 505–511, 2019, doi: 10.1093/jat/bkz014.

[6] T. Buonasera et al., “A comparison of proteomic, genomic, and osteological methods of archaeological sex estimation,” Sci. Rep., vol. 10, no. 1, pp. 1–15, 2020, doi: 10.1038/s41598-020-68550-w.

[7] L. Eckhart, S. Lippens, E. Tschachler, and W. Declercq, “Cell death by cornification,” Biochim. Biophys. Acta - Mol. Cell Res., vol. 1833, no. 12, pp. 3471–3480, 2013, doi: 10.1016/j.bbamcr.2013.06.010.

[8] M. E. Allentoft et al., “The half-life of DNA in bone: Measuring decay kinetics in 158 dated fossils,” Proc. R. Soc. B Biol. Sci., vol. 279, no. 1748, pp. 4724–4733, 2012, doi: 10.1098/rspb.2012.1745.

[9] M. M. Holland et al., “Mitochondrial DNA Sequence Analysis of Human Skeletal Remains: Identification of Remains from the Vietnam War,” J. Forensic Sci., vol. 38, no. 3, p. 13439J, 1993, doi: 10.1520/jfs13439j.

[10] C. Ginther, L. Issel-Tarver, and M. C. King, “Identifying individuals by sequencing mitochondrial DNA from teeth,” Nat. Genet., vol. 2, no. 2, pp. 135–138, 1992, doi: 10.1038/ng1092-135.

[11] D. Fenyö and R. C. Beavis, “A method for assessing the statistical significance of mass spectrometry-based protein identifications using general scoring schemes,” Anal. Chem., vol. 75, no. 4, pp. 768–774, 2003, doi: 10.1021/ac0258709.

[12] D. Fenyö, J. Eriksson, and R. Beavis, “Mass Spectrometric Protein Identification Using the Global Proteome Machine,” in Methods in Molecular Biology, no. 10, 2010, pp. 189–202.

[13] F. Meier et al., “Online parallel accumulation–serial fragmentation (PASEF) with a novel trapped ion mobility mass spectrometer,” Mol. Cell. Proteomics, vol. 17, no. 12, pp. 2534–2545, 2018, doi: 10.1074/mcp.TIR118.000900.

[14] M. Pla-Roca et al., “Antibody colocalization microarray: A scalable technology for multiplex protein analysis in complex samples,” Mol. Cell. Proteomics, vol. 11, no. 4, pp. 1–12, 2012, doi: 10.1074/mcp.M111.011460.

[15] N. Nagaraj et al., “Deep proteome and transcriptome mapping of a human cancer cell line,” Mol. Syst. Biol., vol. 7, no. 548, pp. 1–8, 2011, doi: 10.1038/msb.2011.81.

[16] T. Geiger, A. Wehner, C. Schaab, J. Cox, and M. Mann, “Comparative proteomic analysis of eleven common cell lines reveals ubiquitous but varying expression of most proteins,” Mol. Cell. Proteomics, vol. 11, no. 3, pp. 1–11, 2012, doi: 10.1074/mcp.M111.014050.

[17] Z. C. Goecker, M. R. Salemi, N. Karim, B. S. Phinney, R. H. Rice, and G. J. Parker, “Optimal processing for proteomic genotyping of single human hairs,” Forensic Sci. Int. Genet., vol. 47, no. December 2019, p. 102314, 2020, doi: 10.1016/j.fsigen.2020.102314.

[18] K. E. Mason, P. H. Paul, F. Chu, D. S. Anex, and B. R. Hart, “Development of a Protein-based Human Identification Capability from a Single Hair,” J. Forensic Sci., vol. 64, no. 4, pp. 1152–1159, 2019, doi: 10.1111/1556-4029.13995.

[19] Z. Zhang et al., “Sensitive Method for the Confident Identification of Genetically Variant Peptides in Human Hair Keratin,” J. Forensic Sci., vol. 65, no. 2, pp. 406–420, 2020, doi: 10.1111/1556-4029.14229.

[20] M. J. MacCoss, C. C. Wu, and J. R. Yates, “Probability based validation of protein identifications using a modified SEQUEST algorithm,” Anal. Chem., vol. 74, no. 21, pp. 5593–5599, 2002, doi: 10.1021/ac025826t.

[21] J. Zhang et al., “PEAKS DB: De novo sequencing assisted database search for sensitive and accurate peptide identification,” Mol. Cell. Proteomics, vol. 11, no. 4, pp. 1–8, 2012, doi: 10.1074/mcp.M111.010587.

[22] J. S. Cottrell, “Protein identification using MS/MS data,” J. Proteomics, vol. 74, no. 10, pp. 1842–1851, 2011, doi: 10.1016/j.jprot.2011.05.014.

[23] H. Zhu, S. Pan, S. Gu, E. Morton Bradbury, and X. Chen, “Amino acid residue specific stable isotope labeling for quantitative proteomics,” Rapid Commun. Mass Spectrom., vol. 16, no. 22, pp. 2115–2123, 2002, doi: 10.1002/rcm.831.

[24] Scientific Working Group for Forensic Toxicology (SWGTOX), “Scientific working group for forensic toxicology (SWGTOX) standard practices for method validation in forensic toxicology,” J. Anal. Toxicol., vol. 37, no. 7, pp. 452–474, 2013, doi: 10.1093/jat/bkt054.

[25] European Parliament and the Council of the European Union, “96/23/EC COMMISSION DECISION of 12 August 2002 implementing Council Directive 96/23/EC concerning the performance of analytical methods and the interpretation of results (notified under document number C(2002) 3044)(Text withEEA relevance) (2002/657/EC),” Off. J. Eur. communities, no. L 221/8, pp. 8–36, 2002, doi: 10.1017/CBO9781107415324.004.

[26] World Anti-Doping Agency (WADA), “Identification criteria for qualitative assays incorporating column chromatography and mass spectrometry,” WADA Tech. Doc. - TD2010IDCR, pp. 1–9, 2010, doi: TD2010IDCR.

[27] J. A. Milan et al., “Comparison of protein expression levels and proteomically-inferred genotypes using human hair from different body sites,” Forensic Sci. Int. Genet., vol. 41, no. March, pp. 19–23, 2019, doi: 10.1016/j.fsigen.2019.03.009.

[28] R. N. Franklin, N. Karim, Z. C. Goecker, B. P. Durbin-Johnson, R. H. Rice, and G. J. Parker, “Proteomic genotyping: Using mass spectrometry to infer SNP genotypes in pigmented and non-pigmented hair,” Forensic Sci. Int., vol. 310, 2020, doi: 10.1016/j.forsciint.2020.110200.

[29] S. Gallien, E. Duriez, C. Crone, M. Kellmann, T. Moehring, and B. Domon, “Targeted proteomic quantification on quadrupole-orbitrap mass spectrometer,” Mol. Cell. Proteomics, vol. 11, no. 12, pp. 1709–1723, 2012, doi: 10.1074/mcp.O112.019802.

[30] L. C. Gillet et al., “Targeted data extraction of the MS/MS spectra generated by data-independent acquisition: A new concept for consistent and accurate proteome analysis,” Mol. Cell. Proteomics, vol. 11, no. 6, pp. 1–17, 2012, doi: 10.1074/mcp.O111.016717.

[31] C. Ludwig, L. Gillet, G. Rosenberger, S. Amon, B. C. Collins, and R. Aebersold, “ Data-independent acquisition-based SWATH - MS for quantitative proteomics: a tutorial,” Mol. Syst. Biol., vol. 14, no. 8, pp. 1–23, 2018, doi: 10.15252/msb.20178126.

[32] G. McAlister, S. Eliuk, and R. Huguet, QuanDirect: A simplified approach to fast and accurate, high throughput targeted MS2 quantitation using internal standards..

[33] D. Remane, D. K. Wissenbach, and F. T. Peters, “Recent advances of liquid chromatography–(tandem) mass spectrometry in clinical and forensic toxicology — An update,” Clin. Biochem., vol. 49, no. 13–14, pp. 1051–1071, 2016, doi: 10.1016/j.clinbiochem.2016.07.010.

[34] M. M. Mbughuni, P. J. Jannetto, and L. J. Langman, “Mass spectrometry applications for toxicology,” J. Int. Fed. Clin. Chem. Lab. Med., vol. 27, no. 4, pp. 2016–2043, 2016.

[35] J. Maublanc, S. Dulaurent, J. Morichon, G. Lachâtre, and J. michel Gaulier, “Identification and quantification of 35 psychotropic drugs and metabolites in hair by LC-MS/MS: application in forensic toxicology,” Int. J. Legal Med., vol. 129, no. 2, pp. 259–268, 2015, doi: 10.1007/s00414-014-1005-1.

[36] I. Shah, A. Petroczi, M. Uvacsek, M. Ránky, and D. P. Naughton, “Hair-based rapid analyses for multiple drugs in forensics and doping: Application of dynamic multiple reaction monitoring with LC-MS/MS,” Chem. Cent. J., vol. 8, no. 1, pp. 1–10, 2014, doi: 10.1186/s13065-014-0073-0.

[37] U. Garg and Y. V Zhang, “Mass Spectrometry in Clinical Laboratory: Applications in Therapeutic Drug Monitoring and Toxicology,” Clin. Appl. Mass Spectrom. Drug Anal., vol. 1383, pp. 241–246, 2016, doi: 10.1007/978-1-4939-3252-8.

[38] K. F. Jones, T. L. Carlson, B. A. Eckenrode, and J. Donfack, “Assessing protein sequencing in human single hair shafts of decreasing lengths,” Forensic Sci. Int. Genet., vol. 44, no. September 2019, p. 102145, 2020, doi: 10.1016/j.fsigen.2019.102145.

[39] R. Huguet, S. Eliuk, M. Blank, V. Zabrouskov, and G. McAlister, “A simplified approach to fast and accurate, high throughput targeted MS2 quantitation using internal standard,” 2016.

[40] B. MacLean et al., “Skyline: An open source document editor for creating and analyzing targeted proteomics experiments,” Bioinformatics, vol. 26, no. 7, pp. 966–968, 2010, doi: 10.1093/bioinformatics/btq054.

[41] S. Gessulat et al., “Prosit: proteome-wide prediction of peptide tandem mass spectra by deep learning,” Nat. Methods, vol. 16, no. 6, pp. 509–518, 2019, doi: 10.1038/s41592-019-0426-7.

[42] I. W. Evett and B. Weir, Interpreting DNA evidence: statistical genetics for forensic scientists. Sunderland MA: Sinauer Associates Sunderland MA, 1998.

[43] A. Auton et al., “A global reference for human genetic variation,” Nature, vol. 526, no. 7571, pp. 68–74, 2015, doi: 10.1038/nature15393.

[44] N. Metropolis, A. W. Rosenbluth, M. N. Rosenbluth, A. H. Teller, and E. Teller, “Equation of State Calculations by Fast Computing Machines,” J. Chem. Phys., vol. 21, no. 6, pp. 1087–1092, 1953.

[45] W. K. Hastings, “Monte carlo sampling methods using Markov chains and their applications,” Biometrika, vol. 57, no. 1, pp. 97–109, 1970, doi: 10.1093/biomet/57.1.97.

[46] Y. Perez-Riverol et al., “PRIDE inspector toolsuite: Moving toward a universal visualization tool for proteomics data standard formats and quality assessment of proteomexchange datasets,” Mol. Cell. Proteomics, vol. 15, no. 1, pp. 305–317, 2016, doi: 10.1074/mcp.O115.050229.

[47] V. Sharma et al., “Panorama: A targeted proteomics knowledge base,” J. Proteome Res., vol. 13, no. 9, pp. 4205–4210, 2014, doi: 10.1021/pr5006636.

[48] Y. J. Lee, R. H. Rice, and Y. M. Lee, “Proteome analysis of human hair shaft: From protein identification to posttranslational modification,” Mol. Cell. Proteomics, vol. 5, no. 5, pp. 789–800, 2006, doi: 10.1074/mcp.M500278-MCP200.

[49] L. D. Fricker, “Limitations of Mass Spectrometry-Based Peptidomic Approaches,” J. Am. Soc. Mass Spectrom., vol. 26, no. 12, pp. 1981–1991, 2015, doi: 10.1007/s13361-015-1231-x.

[50] S. R. Wilson, T. Vehus, H. S. Berg, and E. Lundanes, “Nano-LC in proteomics: Recent advances and approaches,” Bioanalysis, vol. 7, no. 14, pp. 1799–1815, 2015, doi: 10.4155/bio.15.92.

[51] H. Henry, H. R. Sobhi, O. Scheibner, M. Bromirski, S. B. Nimkar, and B. Rochat, “Comparison between a high-resolution single-stage Orbitrap and a triple quadrupole mass spectrometer for quantitative analyses of drugs,” Rapid Commun. Mass Spectrom., vol. 26, no. 5, pp. 499–509, 2012, doi: 10.1002/rcm.6121.

[52] E. V Moskovets and A. R. Ivanov, “Comparative studies of peak intensities and chromatographic separation of proteolytic digests, PTMs, and intact proteins obtained by nanoLC-ESI MS analysis at room and elevated temperatures,” Anal. Bioanal. Chem., pp. 3953–3968, 2016, doi: 10.1007/s00216-016-9386-2.

[53] J. C. Gesquiere, E. Diesis, M. T. Cung, and A. Tartar, “Slow isomerization of some proline-containing peptides inducing peak splitting during reversed-phase high-performance liquid chromatography,” J. Chromatogr. A, vol. 478, no. C, pp. 121–129, 1989, doi: 10.1016/0021-9673(89)90010-1.

[54] F. Zhang, W. Ge, G. Ruan, X. Cai, and T. Guo, “Data-Independent Acquisition Mass Spectrometry-Based Proteomics and Software Tools: A Glimpse in 2020,” Proteomics, vol. 1900276, pp. 1–12, 2020, doi: 10.1002/pmic.201900276.

[55] A. Wolf-Yadlin, A. Hu, and W. S. Noble, “Technical advances in proteomics: New developments in data-independent acquisition,” F1000Research, vol. 5, no. 0, pp. 1–12, 2016, doi: 10.12688/f1000research.7042.1.

[56] L. K. Pino, S. C. Just, M. J. MacCoss, and B. C. Searle, “Acquiring and Analyzing Data Independent Acquisition Proteomics Experiments without Spectrum Libraries,” Mol. Cell. Proteomics, vol. 19, no. 7, pp. 1088–1103, 2020, doi: 10.1074/mcp.P119.001913.

[57] L. Stopfer et al., “High-density, targeted monitoring of tyrosine phosphorylation reveals activated signaling networks in human tumors,” bioRxiv, 2020, doi: 10.1101/2020.06.01.127787.

[58] L. M. Smith and N. L. Kelleher, “Proteoform: A single term describing protein complexity,” Nat. Methods, vol. 10, no. 3, pp. 186–187, 2013, doi: 10.1038/nmeth.2369.

[59] G. Rosenberger et al., “Inference and quantification of peptidoforms in large sample cohorts by SWATH-MS,” Nat. Biotechnol., vol. 35, no. 8, pp. 781–788, 2017, doi: 10.1038/nbt.3908.

[60] M. G. Farrell, “Daubert v Merrell Dow Pharmaceutircals, Inc: Epistemiology and Legal Process,” Cardozo Law Rev., vol. 15, pp. 2183–2217, 2014.

[61] Scientific Working Group on DNA Analysis Methods (SWGDAM), “Validation Guidelines for DNA Analysis Methods,” no. December 2016. pp. 1–13, 2016, [Online]. Available: www.swgdam.org.

[62] T. Borja et al., “Proteomic genotyping of fingermark donors with genetically variant peptides,” Forensic Sci. Int. Genet., vol. 42, no. March, pp. 21–30, 2019, doi: 10.1016/j.fsigen.2019.05.005.

[63] G. Parker et al., “Proteomic genotyping: Using mass spectrometry to infer SNP genotypes in a forensic context,” Forensic Sci. Int. Genet. Suppl. Ser., vol. 7, no. 1, pp. 664–666, 2019, doi: 10.1016/j.fsigss.2019.10.130.

[64] Z. C. Goecker, “Forensic proteomics: extracting identifying information from problematic evidence types,” 2019.

[65] L. A. Catlin et al., “Demonstration of a mitochondrial DNA-compatible workflow for genetically variant peptide identification from human hair samples,” Forensic Sci. Int. Genet., vol. 43, no. June, 2019, doi: 10.1016/j.fsigen.2019.102148.

