## Supplemental Figures for "Alternative LC-MS/MS Platforms and Data Acquisition Strategies for Proteomic Genotyping of Human Hair Shafts"

| | | DSQECILTETEAR | | | DSQECILMETEAR | | | GILIDTSR | | | GILVDTSR | | |
|---|---|---|---|---|---|---|---|---|---|---|---|---|---|
| Heavy | Replicate | y7 - 829.4289+ | y6 - 716.3449+ | b4 - 460.1674+ | y7 - 859.4217+ | y6 - 746.3377+ | b4 - 460.1674+ | y6 - 714.4020+ | y5 - 601.3179+ | b3 - 284.1969+ | y5 - 587.3023+ | y4 - 488.2339+ | b3 - 284.1969+ |
| | ZG182 | 92.1% | 100.0% | 37.9% | 100.0% | 82.9% | 34.4% | 72.0% | 100.0% | 52.9% | 100.0% | 85.3% | 44.2% |
| | ZG183 | 100.0% | 96.6% | 37.6% | 100.0% | 83.4% | 34.5% | 70.8% | 100.0% | 51.9% | 100.0% | 83.0% | 44.5% |
| | ZG184 | 100.0% | 91.0% | 34.8% | 100.0% | 82.4% | 35.8% | 74.4% | 100.0% | 52.8% | 100.0% | 83.4% | 43.6% |
| | ZG185 | 100.0% | 91.8% | 36.5% | 100.0% | 82.8% | 34.2% | 68.2% | 100.0% | 49.7% | 100.0% | 86.1% | 46.1% |
| | ZG186 | 100.0% | 98.0% | 35.1% | 100.0% | 83.2% | 34.7% | 72.2% | 100.0% | 52.5% | 100.0% | 84.6% | 44.9% |
| | ZG187 | 100.0% | 98.6% | 35.0% | 100.0% | 80.5% | 33.9% | 69.7% | 100.0% | 49.9% | 100.0% | 84.3% | 43.8% |
| | ZG188 | 100.0% | 95.6% | 35.5% | 100.0% | 82.7% | 34.5% | 70.5% | 100.0% | 49.3% | 100.0% | 85.7% | 44.8% |
| | ZG189 | 99.0% | 100.0% | 39.1% | 100.0% | 83.7% | 34.6% | 68.8% | 100.0% | 51.9% | 100.0% | 84.8% | 46.1% |
| | ZG190 | 100.0% | 92.5% | 32.9% | 100.0% | 84.1% | 35.4% | 74.6% | 100.0% | 56.9% | 100.0% | 84.9% | 44.5% |
| | ZG194 | 98.6% | 100.0% | 38.2% | 100.0% | 82.7% | 34.5% | 64.5% | 100.0% | 51.5% | 100.0% | 84.1% | 44.1% |
| | ZG195 | 100.0% | 86.6% | 33.9% | 100.0% | 84.4% | 34.3% | 70.7% | 100.0% | 54.1% | 100.0% | 83.9% | 44.3% |
| | ZG196 | 100.0% | 94.2% | 34.3% | 100.0% | 82.3% | 33.5% | 70.2% | 100.0% | 51.7% | 100.0% | 82.9% | 43.9% |
| | ZG197 | 98.4% | 100.0% | 35.8% | 100.0% | 80.3% | 34.5% | 77.3% | 100.0% | 54.7% | 100.0% | 83.4% | 44.6% |
| | ZG198 | 100.0% | 99.9% | 37.5% | 100.0% | 83.1% | 34.2% | 70.7% | 100.0% | 52.4% | 100.0% | 85.4% | 44.3% |
| | ZG199 | 100.0% | 92.1% | 32.8% | 100.0% | 80.3% | 33.1% | 73.3% | 100.0% | 55.5% | 100.0% | 85.2% | 44.9% |
| | Average | 99.2% | 95.8% | 35.8% | 100.0% | 82.6% | 34.4% | 71.2% | 100.0% | 52.5% | 100.0% | 84.5% | 44.6% |
| | STD DEV | 2.1% | 4.2% | 2.0% | 0.0% | 1.3% | 0.6% | 3.0% | 0.0% | 2.1% | 0.0% | 1.0% | 0.7% |
| Light | Replicate | y7 - 819.4207+ | y6 - 706.3366+ | b4 - 460.1674+ | y7 - 849.4135+ | y6 - 736.3294+ | b4 - 460.1674+ | y6 - 704.3937+ | y5 - 591.3097+ | b3 - 284.1969+ | y5 - 577.2940+ | y4 - 478.2256+ | b3 - 284.1969+ |
| | ZG182 | 100.0% | 97.1% | 40.3% | 100.0% | 24.7% | 69.1% | 66.5% | 100.0% | 51.1% | 16.0% | 58.9% | 100.0% |
| | ZG183 | 100.0% | 98.9% | 41.2% | 100.0% | 74.8% | 88.5% | 71.8% | 100.0% | 50.6% | 2.8% | 14.4% | 100.0% |
| | ZG184 | 97.0% | 100.0% | 41.8% | 100.0% | 22.5% | 83.2% | 73.0% | 100.0% | 60.7% | 11.3% | 13.9% | 100.0% |
| | ZG185 | 99.6% | 100.0% | 43.6% | 100.0% | 69.0% | 40.7% | 65.7% | 100.0% | 62.2% | 100.0% | 88.1% | 75.7% |
| | ZG186 | 100.0% | 89.5% | 40.3% | 100.0% | 96.6% | 71.3% | 63.6% | 100.0% | 41.7% | 100.0% | 98.7% | 78.1% |
| | ZG187 | 90.0% | 100.0% | 35.5% | 100.0% | 71.9% | 55.7% | 63.7% | 100.0% | 44.7% | 100.0% | 79.8% | 60.6% |
| | ZG188 | 100.0% | 98.5% | 37.6% | 16.7% | 1.1% | 100.0% | 63.7% | 100.0% | 44.0% | 100.0% | 95.3% | 61.8% |
| | ZG189 | 99.6% | 100.0% | 36.7% | 54.3% | 31.6% | 100.0% | 64.8% | 100.0% | 32.2% | 97.7% | 100.0% | 71.3% |
| | ZG190 | 100.0% | 94.6% | 35.7% | 70.6% | 35.2% | 100.0% | 64.3% | 100.0% | 53.6% | 100.0% | 87.5% | 53.6% |
| | ZG194 | 97.4% | 100.0% | 40.1% | 100.0% | 75.9% | 64.7% | 88.3% | 100.0% | 67.4% | 63.9% | 100.0% | 94.4% |
| | ZG195 | 89.9% | 100.0% | 36.3% | 100.0% | 12.7% | 36.0% | 63.7% | 100.0% | 57.1% | 42.3% | 55.8% | 100.0% |
| | ZG196 | 92.4% | 100.0% | 41.7% | 100.0% | 31.8% | 57.2% | 93.6% | 100.0% | 54.2% | 29.3% | 54.6% | 100.0% |
| | ZG197 | 16.5% | 7.6% | 100.0% | 100.0% | 88.2% | 58.0% | 86.9% | 100.0% | 58.6% | 25.8% | 100.0% | 27.0% |
| | ZG198 | 33.5% | 100.0% | 22.7% | 100.0% | 87.1% | 45.7% | 60.2% | 88.4% | 100.0% | 21.2% | 100.0% | 39.4% |
| | ZG199 | 59.7% | 31.0% | 100.0% | 100.0% | 85.6% | 45.2% | 49.0% | 100.0% | 50.7% | 29.7% | 100.0% | 76.3% |

| | | ALETVQER | | | ALETLQER | | | LEGEINTYR | | | LEGEINMYR | | |
|---|---|---|---|---|---|---|---|---|---|---|---|---|---|
| Heavy | Replicate | y5 - 642.3445+ | y4 - 541.2968+ | b3 - 314.1710+ | y5 - 656.3601+ | y4 - 555.3125+ | b3 - 314.1710+ | y8 - 991.4719+ | y7 - 862.4293+ | b2 - 243.1339+ | y8 - 1021.4647+ | y7 - 892.4221+ | b2 - 243.1339+ |
| | ZG182 | 100.0% | 38.2% | 36.9% | 100.0% | 41.7% | 43.4% | 18.0% | 100.0% | 94.5% | 15.4% | 100.0% | 88.6% |
| | ZG183 | 100.0% | 40.7% | 36.2% | 100.0% | 42.0% | 47.0% | 17.5% | 100.0% | 99.9% | 16.7% | 100.0% | 89.6% |
| | ZG184 | 100.0% | 37.1% | 40.5% | 100.0% | 44.5% | 49.1% | 18.3% | 100.0% | 97.1% | 15.6% | 100.0% | 85.1% |
| | ZG185 | 100.0% | 37.4% | 39.9% | 100.0% | 40.6% | 43.8% | 19.0% | 100.0% | 96.4% | 15.7% | 100.0% | 93.6% |
| | ZG186 | 100.0% | 37.4% | 38.2% | 100.0% | 39.5% | 41.8% | 18.6% | 100.0% | 91.8% | 16.1% | 100.0% | 88.3% |
| | ZG187 | 100.0% | 40.8% | 38.8% | 100.0% | 43.0% | 44.8% | 19.1% | 100.0% | 93.5% | 16.9% | 100.0% | 97.1% |
| | ZG188 | 100.0% | 37.6% | 39.0% | 100.0% | 43.2% | 42.4% | 18.1% | 100.0% | 92.6% | 15.8% | 100.0% | 90.8% |
| | ZG189 | 100.0% | 38.1% | 37.3% | 100.0% | 42.2% | 44.4% | 18.7% | 100.0% | 92.3% | 16.4% | 100.0% | 92.3% |
| | ZG190 | 100.0% | 36.0% | 37.0% | 100.0% | 43.9% | 46.1% | 19.2% | 100.0% | 94.9% | 17.3% | 98.8% | 100.0% |
| | ZG194 | 100.0% | 37.0% | 36.1% | 100.0% | 41.2% | 44.4% | 17.5% | 100.0% | 87.1% | 16.2% | 100.0% | 93.6% |
| | ZG195 | 100.0% | 38.3% | 37.7% | 100.0% | 42.6% | 42.5% | 17.2% | 100.0% | 92.7% | 16.0% | 100.0% | 88.1% |
| | ZG196 | 100.0% | 36.1% | 37.5% | 100.0% | 40.4% | 41.8% | 18.3% | 100.0% | 92.8% | 17.8% | 100.0% | 93.8% |
| | ZG197 | 100.0% | 36.4% | 35.5% | 100.0% | 42.1% | 44.3% | 18.1% | 100.0% | 94.5% | 17.4% | 100.0% | 96.6% |
| | ZG198 | 100.0% | 37.9% | 35.4% | 100.0% | 44.5% | 44.3% | 17.9% | 100.0% | 95.6% | 15.9% | 100.0% | 86.0% |
| | ZG199 | 100.0% | 37.2% | 36.7% | 100.0% | 38.8% | 40.3% | 17.8% | 100.0% | 89.4% | 16.3% | 100.0% | 90.8% |
| | Average | 100.0% | 37.7% | 37.5% | 100.0% | 42.0% | 44.0% | 18.2% | 100.0% | 93.7% | 16.4% | 99.9% | 91.6% |
| | STD DEV | 0.0% | 1.4% | 1.5% | 0.0% | 1.7% | 2.2% | 0.6% | 0.0% | 3.1% | 0.7% | 0.3% | 4.2% |
| Light | Replicate | y5 - 632.3362+ | y4 - 531.2885+ | b3 - 314.1710+ | y5 - 646.3519+ | y4 - 545.3042+ | b3 - 314.1710+ | y8 - 981.4636+ | y7 - 852.4210+ | b2 - 243.1339+ | y8 - 1011.4564+ | y7 - 882.4138+ | b2 - 243.1339+ |
| | ZG182 | 100.0% | 63.3% | 94.6% | 100.0% | 34.4% | 2.7% | 18.9% | 100.0% | 97.6% | 21.4% | 100.0% | 92.0% |
| | ZG183 | 33.7% | 75.2% | 100.0% | 14.0% | 100.0% | 1.5% | 18.5% | 97.8% | 100.0% | 18.0% | 100.0% | 92.1% |
| | ZG184 | 37.3% | 100.0% | 51.2% | 27.0% | 100.0% | 38.1% | 19.0% | 95.7% | 100.0% | 21.4% | 100.0% | 88.5% |
| | ZG185 | 70.6% | 59.5% | 100.0% | 100.0% | 52.1% | 44.2% | 18.5% | 100.0% | 97.0% | 14.7% | 100.0% | 91.9% |
| | ZG186 | 53.9% | 18.2% | 100.0% | 100.0% | 58.5% | 23.6% | 18.9% | 99.1% | 100.0% | 28.9% | 100.0% | 93.8% |
| | ZG187 | 43.8% | 25.3% | 100.0% | 100.0% | 52.1% | 21.5% | 18.8% | 100.0% | 98.0% | 17.7% | 100.0% | 78.8% |
| | ZG188 | 100.0% | 49.0% | 46.5% | 56.8% | 100.0% | 8.1% | 18.4% | 100.0% | 99.1% | 17.5% | 35.4% | 100.0% |
| | ZG189 | 100.0% | 37.4% | 28.6% | 100.0% | 91.1% | 73.8% | 19.0% | 100.0% | 99.6% | 27.0% | 100.0% | 53.1% |
| | ZG190 | 100.0% | 22.0% | 23.2% | 33.9% | 100.0% | 45.2% | 19.0% | 100.0% | 96.2% | 14.7% | 12.2% | 100.0% |
| | ZG194 | 82.7% | 100.0% | 57.6% | 100.0% | 49.7% | 15.3% | 19.1% | 100.0% | 98.0% | 21.6% | 100.0% | 13.8% |
| | ZG195 | 96.9% | 100.0% | 72.6% | 100.0% | 49.6% | 26.8% | 18.9% | 100.0% | 96.0% | 25.6% | 85.5% | 100.0% |
| | ZG196 | 91.1% | 21.9% | 100.0% | 100.0% | 45.5% | 26.3% | 19.5% | 99.9% | 100.0% | 13.0% | 100.0% | 57.3% |
| | ZG197 | 4.8% | 100.0% | 32.3% | 43.9% | 97.8% | 100.0% | 19.4% | 100.0% | 95.4% | 23.8% | 100.0% | 21.4% |
| | ZG198 | 21.8% | 100.0% | 96.2% | 99.4% | 100.0% | 82.0% | 19.9% | 100.0% | 97.7% | 19.4% | 100.0% | 95.3% |
| | ZG199 | 45.7% | 100.0% | 89.2% | 62.0% | 100.0% | 34.6% | 18.7% | 100.0% | 97.6% | 17.9% | 100.0% | 92.0% |

| | | YISLIYTNYEAGKDDYVK | | | YVSLIYTNYEAGKDDYVK | | | VSAMYSSSSCKLPSLSPVAR | | | VSAMYSSSPCKLPSLSPVAR | | |
|---|---|---|---|---|---|---|---|---|---|---|---|---|---|
| Heavy | Replicate | y8 - 903.4662+ | b2 - 277.1547+ | b3 - 364.1867+ | y12 - 1410.6627+ | b2 - 263.1390+ | b3 - 350.1710+ | y8 - 836.4864+ | y5 - 539.3175+ | b3 - 258.1448+ | y8 - 836.4864+ | y5 - 539.3175+ | b3 - 258.1448+ |
| | ZG182 | 2.8% | 100.0% | 18.5% | 92.2% | 100.0% | 75.4% | 100.0% | 29.5% | 30.7% | 100.0% | 30.3% | 27.1% |
| | ZG183 | 2.8% | 100.0% | 18.1% | 87.9% | 100.0% | 75.7% | 100.0% | 30.9% | 33.6% | 100.0% | 27.7% | 25.4% |
| | ZG184 | 3.2% | 100.0% | 19.0% | 89.3% | 100.0% | 77.3% | 100.0% | 31.2% | 31.8% | 100.0% | 28.6% | 26.3% |
| | ZG185 | 3.0% | 100.0% | 18.0% | 94.4% | 100.0% | 91.7% | 100.0% | 28.9% | 31.0% | 100.0% | 29.9% | 28.8% |
| | ZG186 | 3.1% | 100.0% | 19.5% | 85.9% | 100.0% | 78.7% | 100.0% | 30.0% | 31.9% | 100.0% | 29.7% | 27.3% |
| | ZG187 | 3.1% | 100.0% | 20.3% | 98.3% | 100.0% | 82.0% | 100.0% | 31.4% | 32.2% | 100.0% | 28.8% | 26.2% |
| | ZG188 | 3.3% | 100.0% | 19.1% | 88.5% | 100.0% | 76.1% | 100.0% | 31.5% | 34.2% | 100.0% | 28.6% | 25.3% |
| | ZG189 | 3.0% | 100.0% | 19.5% | 82.3% | 100.0% | 70.8% | 100.0% | 30.3% | 31.3% | 100.0% | 30.0% | 26.1% |
| | ZG190 | 3.1% | 100.0% | 19.0% | 84.5% | 100.0% | 73.7% | 100.0% | 31.5% | 32.7% | 100.0% | 33.5% | 29.1% |
| | ZG194 | 3.5% | 100.0% | 20.1% | 85.4% | 100.0% | 75.1% | 100.0% | 32.1% | 33.2% | 100.0% | 29.2% | 25.8% |
| | ZG195 | 3.1% | 100.0% | 18.7% | 87.4% | 100.0% | 77.8% | 100.0% | 30.9% | 30.6% | 100.0% | 29.8% | 26.0% |
| | ZG196 | 3.0% | 100.0% | 19.4% | 87.8% | 100.0% | 77.9% | 100.0% | 32.7% | 32.9% | 100.0% | 30.1% | 25.3% |
| | ZG197 | 3.1% | 100.0% | 19.3% | 91.2% | 100.0% | 77.1% | 100.0% | 29.8% | 30.7% | 100.0% | 28.6% | 25.8% |
| | ZG198 | 3.1% | 100.0% | 19.6% | 81.0% | 100.0% | 75.6% | 100.0% | 31.5% | 31.6% | 100.0% | 30.0% | 27.1% |
| | ZG199 | 3.2% | 100.0% | 18.8% | 91.6% | 100.0% | 83.3% | 100.0% | 27.8% | 30.5% | 100.0% | 29.2% | 26.7% |
| | Average | 3.1% | 100.0% | 19.1% | 88.5% | 100.0% | 77.9% | 100.0% | 30.7% | 31.9% | 100.0% | 29.6% | 26.6% |
| | STD DEV | 0.2% | 0.0% | 0.6% | 4.6% | 0.0% | 4.9% | 0.0% | 1.3% | 1.2% | 0.0% | 1.3% | 1.2% |
| Light | Replicate | y8 - 895.4520+ | b2 - 277.1547+ | b3 - 364.1867+ | y12 - 1402.6485+ | b2 - 263.1390+ | b3 - 350.1710+ | y8 - 826.4781+ | y5 - 529.3093+ | b3 - 258.1448+ | y8 - 826.4781+ | y5 - 529.3093+ | b3 - 258.1448+ |
| | ZG182 | 4.0% | 100.0% | 20.6% | 23.8% | 100.0% | 17.4% | 100.0% | 73.2% | 46.9% | 0.3% | 0.2% | 100.0% |
| | ZG183 | 2.5% | 100.0% | 16.7% | 9.8% | 96.7% | 100.0% | 100.0% | 29.3% | 13.3% | 0.9% | 0.6% | 100.0% |
| | ZG184 | 6.2% | 100.0% | 21.3% | 47.4% | 100.0% | 80.3% | 100.0% | 15.8% | 28.4% | 1.9% | 3.3% | 100.0% |
| | ZG185 | 6.3% | 100.0% | 23.4% | 100.0% | 61.2% | 16.0% | 63.6% | 20.2% | 100.0% | 1.0% | 0.6% | 100.0% |
| | ZG186 | 7.1% | 100.0% | 18.2% | 67.4% | 100.0% | 38.8% | 57.6% | 61.3% | 100.0% | 1.4% | 0.9% | 100.0% |
| | ZG187 | 6.1% | 100.0% | 21.7% | 100.0% | 94.7% | 58.5% | 77.2% | 18.5% | 100.0% | 3.5% | 0.4% | 100.0% |
| | ZG188 | 6.0% | 100.0% | 14.5% | 70.8% | 57.1% | 100.0% | 100.0% | 2.6% | 33.5% | 1.4% | 2.4% | 100.0% |
| | ZG189 | 9.6% | 100.0% | 18.6% | 100.0% | 95.1% | 55.4% | 60.7% | 5.9% | 100.0% | 1.4% | 0.1% | 100.0% |
| | ZG190 | 6.6% | 100.0% | 25.4% | 100.0% | 45.7% | 31.9% | 100.0% | 50.9% | 49.8% | 1.1% | 0.9% | 100.0% |
| | ZG194 | 51.7% | 98.0% | 100.0% | 100.0% | 92.2% | 46.9% | 100.0% | 19.3% | 19.5% | 1.6% | 0.2% | 100.0% |
| | ZG195 | 16.9% | 100.0% | 28.6% | 100.0% | 27.4% | 28.5% | 3.6% | 33.8% | 100.0% | 0.6% | 0.2% | 100.0% |
| | ZG196 | 40.9% | 100.0% | 42.5% | 29.7% | 100.0% | 23.6% | 81.7% | 11.7% | 100.0% | 2.1% | 1.2% | 100.0% |
| | ZG197 | 11.7% | 100.0% | 27.0% | 81.5% | 100.0% | 66.8% | 100.0% | 16.5% | 28.4% | 0.2% | 1.0% | 100.0% |
| | ZG198 | 7.9% | 100.0% | 18.2% | 100.0% | 49.9% | 32.7% | 62.6% | 37.3% | 100.0% | 2.0% | 0.0% | 100.0% |
| | ZG199 | 3.0% | 100.0% | 21.2% | 100.0% | 70.0% | 23.7% | 100.0% | 56.9% | 22.6% | 0.9% | 1.6% | 100.0% |

| | | EHCSACGPLSR | | | EHCSACGPLSQLLVK | | | EWSTFAVGPGHCLQLNDR | | | EWSTFAVGPGHCLQLHDR | | |
| --- | --- | --- | --- | --- | --- | --- | --- | --- | --- | --- | --- | --- | --- |
| Heavy | Replicate | y9 - 1017.4480+ | y8 - 857.4173+ | b3 - 427.1394+ | y13 - 1440.7429+ | y9 - 962.6124+ | b3 - 427.1394+ | y5 - 655.3397+ | y4 - 527.2812+ | b6 - 722.3144+ | y11 - 1299.6250+ | y3 - 437.2131+ | b6 - 722.3144+ |
| | ZG182 | 100.0% | 47.3% | 29.3% | 100.0% | 90.6% | 79.5% | 85.5% | 100.0% | 29.5% | 45.7% | 100.0% | 61.0% |
| | ZG183 | 100.0% | 47.1% | 28.5% | 100.0% | 83.2% | 90.8% | 78.9% | 100.0% | 26.5% | 41.6% | 100.0% | 51.6% |
| | ZG184 | 100.0% | 47.3% | 27.0% | 100.0% | 91.3% | 93.3% | 75.9% | 100.0% | 26.7% | 44.5% | 100.0% | 60.4% |
| | ZG185 | 100.0% | 50.0% | 29.5% | 94.8% | 100.0% | 90.3% | 84.5% | 100.0% | 29.8% | 43.1% | 100.0% | 49.4% |
| | ZG186 | 100.0% | 46.9% | 29.0% | 100.0% | 88.5% | 81.3% | 85.1% | 100.0% | 30.0% | 43.8% | 100.0% | 52.9% |
| | ZG187 | 100.0% | 45.3% | 25.5% | 100.0% | 82.2% | 89.0% | 78.7% | 100.0% | 26.6% | 39.3% | 100.0% | 50.0% |
| | ZG188 | 100.0% | 45.2% | 27.5% | 100.0% | 95.7% | 92.9% | 78.8% | 100.0% | 27.4% | 45.3% | 100.0% | 51.1% |
| | ZG189 | 100.0% | 51.7% | 29.8% | 100.0% | 89.7% | 95.6% | 81.0% | 100.0% | 30.3% | 42.9% | 100.0% | 54.1% |
| | ZG190 | 100.0% | 48.3% | 26.9% | 100.0% | 86.2% | 79.9% | 79.3% | 100.0% | 28.6% | 46.9% | 100.0% | 56.4% |
| | ZG194 | 100.0% | 47.6% | 27.8% | 100.0% | 85.2% | 81.5% | 81.6% | 100.0% | 29.1% | 46.5% | 100.0% | 52.0% |
| | ZG195 | 100.0% | 45.9% | 27.5% | 100.0% | 88.3% | 97.2% | 81.3% | 100.0% | 27.4% | 47.5% | 100.0% | 49.3% |
| | ZG196 | 100.0% | 49.0% | 27.4% | 100.0% | 94.4% | 93.4% | 84.6% | 100.0% | 29.2% | 45.8% | 100.0% | 50.9% |
| | ZG197 | 100.0% | 47.1% | 26.5% | 100.0% | 87.4% | 92.1% | 78.5% | 100.0% | 28.8% | 45.6% | 100.0% | 57.2% |
| | ZG198 | 100.0% | 44.4% | 27.3% | 100.0% | 97.2% | 89.7% | 82.6% | 100.0% | 29.8% | 48.4% | 100.0% | 52.4% |
| | ZG199 | 100.0% | 46.8% | 27.1% | 100.0% | 85.5% | 92.2% | 80.0% | 100.0% | 28.8% | 44.7% | 100.0% | 53.8% |
| | Average | 100.0% | 47.3% | 27.8% | 99.7% | 89.7% | 89.2% | 81.1% | 100.0% | 28.6% | 44.8% | 100.0% | 53.5% |
| | STD DEV | 0.0% | 1.9% | 1.2% | 1.3% | 5.2% | 5.8% | 2.9% | 0.0% | 1.3% | 2.4% | 0.0% | 3.7% |
| Light | Replicate | y9 - 1007.4397+ | y8 - 847.4091+ | b3 - 427.1394+ | y13 - 1432.7287+ | y9 - 954.5982+ | b3 - 427.1394+ | y5 - 645.3315+ | y4 - 517.2729+ | b6 - 722.3144+ | y11 - 1289.6168+ | y3 - 427.2048+ | b6 - 722.3144+ |
| | ZG182 | 55.5% | 24.6% | 100.0% | 15.0% | 39.0% | 100.0% | 59.8% | 100.0% | 74.9% | 6.8% | 54.8% | 100.0% |
| | ZG183 | 56.3% | 25.7% | 100.0% | 10.8% | 100.0% | 68.7% | 90.5% | 100.0% | 37.6% | 40.0% | 3.0% | 100.0% |
| | ZG184 | 53.7% | 24.3% | 100.0% | 3.4% | 100.0% | 56.9% | 100.0% | 81.6% | 51.8% | 18.0% | 90.3% | 100.0% |
| | ZG185 | 75.4% | 37.7% | 100.0% | 27.1% | 22.2% | 100.0% | 100.0% | 59.0% | 18.6% | 7.2% | 11.7% | 100.0% |
| | ZG186 | 80.7% | 36.5% | 100.0% | 19.5% | 100.0% | 83.3% | 100.0% | 27.6% | 29.1% | 12.6% | 34.2% | 100.0% |
| | ZG187 | 94.3% | 44.2% | 100.0% | 16.6% | 100.0% | 47.9% | 100.0% | 38.9% | 32.1% | 72.0% | 13.5% | 100.0% |
| | ZG188 | 69.0% | 32.1% | 100.0% | 16.3% | 64.9% | 100.0% | 57.5% | 80.4% | 100.0% | 4.9% | 9.2% | 100.0% |
| | ZG189 | 96.5% | 43.6% | 100.0% | 20.1% | 88.6% | 100.0% | 1.5% | 46.6% | 100.0% | 52.2% | 58.4% | 100.0% |
| | ZG190 | 54.8% | 25.1% | 100.0% | 38.6% | 6.8% | 100.0% | 100.0% | 31.9% | 88.1% | 32.1% | 49.7% | 100.0% |
| | ZG194 | 27.5% | 12.9% | 100.0% | 24.8% | 56.6% | 100.0% | 100.0% | 43.0% | 46.7% | 20.4% | 53.5% | 100.0% |
| | ZG195 | 24.1% | 12.5% | 100.0% | 17.5% | 100.0% | 83.1% | 10.4% | 1.3% | 100.0% | 15.4% | 5.4% | 100.0% |
| | ZG196 | 26.4% | 10.1% | 100.0% | 14.5% | 76.1% | 100.0% | 56.2% | 100.0% | 69.8% | 16.3% | 22.7% | 100.0% |
| | ZG197 | 100.0% | 37.6% | 28.4% | 6.6% | 100.0% | 31.2% | 100.0% | 9.8% | 23.0% | 36.0% | 81.2% | 100.0% |
| | ZG198 | 100.0% | 37.3% | 30.8% | 18.8% | 79.7% | 100.0% | 80.8% | 82.2% | 100.0% | 61.9% | 100.0% | 35.4% |
| | ZG199 | 100.0% | 46.0% | 32.0% | 28.6% | 88.5% | 100.0% | 100.0% | 68.0% | 25.4% | 100.0% | 50.6% | 10.7% |

| | | GVALSNVIHK | | | GVALSNVVHK | | | DLNMDCMVAEIK | | | DLNMDCIVAEIK | | |
| --- | --- | --- | --- | --- | --- | --- | --- | --- | --- | --- | --- | --- | --- |
| Heavy | Replicate | y6 - 705.4133+ | y3 - 405.2700+ | b3 - 228.1343+ | y8 - 875.5189+ | y7 - 804.4818+ | y6 - 691.3977+ | y8 - 973.4573+ | y6 - 698.3997+ | b3 - 343.1612+ | y8 - 955.5008+ | y6 - 680.4433+ | b3 - 343.1612+ |
| | ZG182 | 71.8% | 10.3% | 100.0% | 76.9% | 41.6% | 100.0% | 100.0% | 16.5% | 86.7% | 100.0% | 16.3% | 81.6% |
| | ZG183 | 77.0% | 10.3% | 100.0% | 70.4% | 40.5% | 100.0% | 100.0% | 16.1% | 87.0% | 100.0% | 15.5% | 81.9% |
| | ZG184 | 71.3% | 10.3% | 100.0% | 72.4% | 39.0% | 100.0% | 100.0% | 16.2% | 86.6% | 100.0% | 15.8% | 84.0% |
| | ZG185 | 76.2% | 10.3% | 100.0% | 73.5% | 40.8% | 100.0% | 100.0% | 16.4% | 84.4% | 100.0% | 16.4% | 83.1% |
| | ZG186 | 70.0% | 9.6% | 100.0% | 73.5% | 39.4% | 100.0% | 100.0% | 15.7% | 82.8% | 100.0% | 16.9% | 82.2% |
| | ZG187 | 72.6% | 9.5% | 100.0% | 73.4% | 39.4% | 100.0% | 100.0% | 15.9% | 83.7% | 100.0% | 16.3% | 82.0% |
| | ZG188 | 72.2% | 9.4% | 100.0% | 72.1% | 39.6% | 100.0% | 100.0% | 16.1% | 85.1% | 100.0% | 16.9% | 81.8% |
| | ZG189 | 72.7% | 9.4% | 100.0% | 71.9% | 39.0% | 100.0% | 100.0% | 16.0% | 87.1% | 100.0% | 16.7% | 85.9% |
| | ZG190 | 75.8% | 9.5% | 100.0% | 69.6% | 40.2% | 100.0% | 100.0% | 15.8% | 84.1% | 100.0% | 16.9% | 86.7% |
| | ZG194 | 72.5% | 9.9% | 100.0% | 74.4% | 39.7% | 100.0% | 100.0% | 16.4% | 88.0% | 100.0% | 15.3% | 83.0% |
| | ZG195 | 79.2% | 10.5% | 100.0% | 71.6% | 39.7% | 100.0% | 100.0% | 16.0% | 84.3% | 100.0% | 16.7% | 87.1% |
| | ZG196 | 71.6% | 9.1% | 100.0% | 70.1% | 39.2% | 100.0% | 100.0% | 15.9% | 83.2% | 100.0% | 16.3% | 78.7% |
| | ZG197 | 67.8% | 10.1% | 100.0% | 71.6% | 39.2% | 100.0% | 100.0% | 16.2% | 86.5% | 100.0% | 16.7% | 86.1% |
| | ZG198 | 73.8% | 10.5% | 100.0% | 70.3% | 39.3% | 100.0% | 100.0% | 16.4% | 83.4% | 100.0% | 17.2% | 81.0% |
| | ZG199 | 69.9% | 9.4% | 100.0% | 75.7% | 40.6% | 100.0% | 100.0% | 16.3% | 87.1% | 100.0% | 15.7% | 82.1% |
| | Average | 73.0% | 9.9% | 100.0% | 72.5% | 39.8% | 100.0% | 100.0% | 16.1% | 85.3% | 100.0% | 16.4% | 83.2% |
| | STD DEV | 3.0% | 0.5% | 0.0% | 2.1% | 0.8% | 0.0% | 0.0% | 0.2% | 1.7% | 0.0% | 0.6% | 2.4% |
| Light | Replicate | y6 - 697.3991+ | y3 - 397.2558+ | b3 - 228.1343+ | y8 - 867.5047+ | y7 - 796.4676+ | y6 - 683.3835+ | y8 - 965.4431+ | y6 - 690.3855+ | b3 - 343.1612+ | y8 - 947.4866+ | y6 - 672.4291+ | b3 - 343.1612+ |
| | ZG182 | 31.6% | 42.7% | 100.0% | 100.0% | 24.4% | 60.3% | 100.0% | 2.3% | 72.6% | 100.0% | 16.8% | 80.2% |
| | ZG183 | 54.2% | 55.3% | 100.0% | 100.0% | 22.9% | 82.1% | 100.0% | 28.7% | 33.2% | 100.0% | 17.5% | 79.7% |
| | ZG184 | 56.7% | 18.6% | 100.0% | 93.0% | 25.9% | 100.0% | 100.0% | 6.5% | 85.1% | 100.0% | 17.9% | 80.7% |
| | ZG185 | 63.0% | 28.4% | 100.0% | 100.0% | 29.2% | 76.5% | 100.0% | 36.2% | 13.3% | 100.0% | 17.4% | 78.0% |
| | ZG186 | 62.1% | 28.5% | 100.0% | 100.0% | 25.8% | 83.5% | 87.2% | 100.0% | 20.9% | 100.0% | 16.9% | 80.4% |
| | ZG187 | 48.3% | 11.8% | 100.0% | 91.5% | 26.4% | 100.0% | 100.0% | 67.0% | 59.6% | 100.0% | 17.2% | 75.6% |
| | ZG188 | 32.2% | 23.8% | 100.0% | 86.2% | 34.2% | 100.0% | 100.0% | 64.6% | 71.2% | 100.0% | 17.0% | 83.0% |
| | ZG189 | 54.2% | 26.6% | 100.0% | 100.0% | 24.8% | 82.4% | 100.0% | 31.1% | 55.0% | 100.0% | 17.1% | 81.8% |
| | ZG190 | 46.2% | 53.5% | 100.0% | 91.2% | 35.8% | 100.0% | 100.0% | 82.5% | 41.7% | 100.0% | 17.7% | 77.4% |
| | ZG194 | 29.9% | 15.1% | 100.0% | 100.0% | 31.9% | 92.3% | 94.7% | 25.2% | 100.0% | 77.4% | 100.0% | 47.4% |
| | ZG195 | 46.8% | 46.3% | 100.0% | 100.0% | 32.9% | 82.1% | 100.0% | 35.2% | 82.4% | 100.0% | 16.4% | 14.7% |
| | ZG196 | 59.8% | 95.5% | 100.0% | 96.9% | 31.6% | 100.0% | 100.0% | 27.5% | 78.2% | 100.0% | 48.7% | 66.4% |
| | ZG197 | 65.3% | 46.3% | 100.0% | 100.0% | 32.2% | 90.2% | 100.0% | 21.1% | 84.9% | 24.7% | 100.0% | 11.2% |
| | ZG198 | 53.6% | 24.3% | 100.0% | 100.0% | 63.1% | 58.3% | 100.0% | 16.0% | 69.3% | 42.0% | 100.0% | 8.8% |
| | ZG199 | 70.2% | 22.0% | 100.0% | 100.0% | 70.0% | 11.6% | 100.0% | 28.3% | 81.3% | 54.8% | 3.7% | 100.0% |

| | | GAFLYEPCGVSTPVLSTGVLR | | | GAFLYDPCGVSTPVLSTGVLR | | | ARPLEQAVAAIVCTFQEYAGR | | | AKPLEQAVAAIVCTFQEYAGR | | |
| --- | --- | --- | --- | --- | --- | --- | --- | --- | --- | --- | --- | --- | --- |
| Heavy | Replicate | y9 - 951.5861+ | b6 - 681.3243+ | b10 - 1094.4975+ | y9 - 951.5861+ | b6 - 667.3086+ | b10 - 1080.4819+ | y13 - 1495.7237+ | b2 - 270.1561+ | b11 - 1162.6579+ | y13 - 1495.7237+ | b2 - 242.1499+ | b11 - 1134.6517+ |
| | ZG182 | 67.0% | 100.0% | 30.7% | 100.0% | 82.2% | 58.2% | 12.6% | 35.4% | 100.0% | 100.0% | 58.6% | 40.8% |
| | ZG183 | 64.3% | 100.0% | 31.3% | 100.0% | 76.8% | 54.6% | 12.2% | 35.7% | 100.0% | 100.0% | 56.2% | 39.3% |
| | ZG184 | 61.5% | 100.0% | 31.4% | 100.0% | 79.1% | 57.0% | 12.3% | 34.5% | 100.0% | 100.0% | 56.3% | 37.8% |
| | ZG185 | 59.8% | 100.0% | 29.8% | 100.0% | 74.0% | 51.6% | 11.9% | 32.5% | 100.0% | 100.0% | 59.3% | 41.3% |
| | ZG186 | 60.5% | 100.0% | 27.0% | 100.0% | 79.0% | 58.6% | 12.1% | 35.6% | 100.0% | 100.0% | 59.5% | 43.5% |
| | ZG187 | 62.8% | 100.0% | 30.4% | 100.0% | 77.2% | 61.9% | 11.9% | 34.5% | 100.0% | 100.0% | 59.1% | 40.8% |
| | ZG188 | 63.4% | 100.0% | 30.3% | 100.0% | 77.3% | 54.5% | 10.7% | 34.0% | 100.0% | 100.0% | 61.6% | 38.9% |
| | ZG189 | 62.7% | 100.0% | 29.9% | 100.0% | 78.9% | 53.8% | 12.9% | 35.4% | 100.0% | 100.0% | 58.8% | 41.2% |
| | ZG190 | 64.6% | 100.0% | 30.1% | 100.0% | 77.6% | 54.1% | 12.6% | 35.1% | 100.0% | 100.0% | 56.6% | 41.6% |
| | ZG194 | 60.5% | 100.0% | 29.7% | 100.0% | 72.9% | 52.6% | 12.2% | 35.7% | 100.0% | 100.0% | 59.3% | 40.4% |
| | ZG195 | 65.2% | 100.0% | 31.6% | 100.0% | 75.6% | 52.2% | 12.4% | 35.9% | 100.0% | 100.0% | 58.7% | 41.1% |
| | ZG196 | 64.4% | 100.0% | 29.3% | 100.0% | 79.2% | 54.2% | 12.8% | 34.9% | 100.0% | 100.0% | 59.7% | 39.7% |
| | ZG197 | 62.6% | 100.0% | 29.5% | 100.0% | 75.3% | 55.6% | 12.3% | 34.5% | 100.0% | 100.0% | 59.0% | 45.0% |
| | ZG198 | 65.9% | 100.0% | 30.4% | 100.0% | 80.6% | 54.5% | 12.8% | 33.9% | 100.0% | 100.0% | 58.6% | 40.8% |
| | ZG199 | 63.3% | 100.0% | 31.0% | 100.0% | 78.9% | 57.3% | 11.8% | 35.0% | 100.0% | 100.0% | 56.2% | 36.7% |
| | Average | 63.2% | 100.0% | 30.2% | 100.0% | 77.6% | 55.4% | 12.2% | 34.8% | 100.0% | 100.0% | 58.5% | 40.6% |
| | STD DEV | 2.1% | 0.0% | 1.1% | 0.0% | 2.5% | 2.8% | 0.6% | 0.9% | 0.0% | 0.0% | 1.5% | 2.0% |
| Light | Replicate | y9 - 941.5778+ | b6 - 681.3243+ | b10 - 1094.4975+ | y9 - 941.5778+ | b6 - 667.3086+ | b10 - 1080.4819+ | y13 - 1485.7155+ | b2 - 270.1561+ | b11 - 1162.6579+ | y13 - 1485.7155+ | b2 - 242.1499+ | b11 - 1134.6517+ |
| | ZG182 | 64.1% | 11.4% | 100.0% | 84.4% | 33.0% | 100.0% | 12.9% | 37.4% | 100.0% | 1.9% | 13.7% | 100.0% |
| | ZG183 | 94.6% | 7.2% | 100.0% | 44.0% | 17.3% | 100.0% | 13.6% | 37.4% | 100.0% | 48.0% | 100.0% | 16.4% |
| | ZG184 | 50.8% | 13.7% | 100.0% | 18.1% | 14.3% | 100.0% | 13.0% | 34.7% | 100.0% | 63.1% | 16.8% | 100.0% |
| | ZG185 | 53.0% | 100.0% | 41.4% | 34.9% | 9.3% | 100.0% | 11.9% | 44.8% | 100.0% | 100.0% | 14.6% | 38.5% |
| | ZG186 | 63.2% | 100.0% | 39.5% | 15.2% | 40.5% | 100.0% | 13.0% | 44.2% | 100.0% | 100.0% | 9.8% | 39.2% |
| | ZG187 | 64.2% | 100.0% | 38.6% | 26.1% | 17.2% | 100.0% | 12.2% | 38.8% | 100.0% | 100.0% | 12.2% | 37.1% |
| | ZG188 | 64.8% | 100.0% | 60.8% | 13.0% | 46.7% | 100.0% | 12.5% | 40.0% | 100.0% | 68.4% | 95.2% | 100.0% |
| | ZG189 | 68.3% | 100.0% | 57.0% | 100.0% | 44.0% | 87.8% | 12.9% | 39.2% | 100.0% | 100.0% | 59.9% | 28.1% |
| | ZG190 | 59.6% | 100.0% | 46.8% | 14.1% | 8.2% | 100.0% | 13.5% | 40.0% | 100.0% | 92.7% | 2.6% | 100.0% |
| | ZG194 | 7.2% | 5.6% | 100.0% | 100.0% | 86.1% | 72.3% | 15.8% | 64.9% | 100.0% | 100.0% | 4.2% | 34.4% |
| | ZG195 | 4.4% | 0.1% | 100.0% | 100.0% | 72.9% | 77.5% | 12.9% | 54.6% | 100.0% | 100.0% | 12.4% | 36.2% |
| | ZG196 | 29.4% | 100.0% | 75.7% | 93.6% | 100.0% | 57.9% | 13.4% | 77.3% | 100.0% | 100.0% | 13.7% | 29.6% |
| | ZG197 | 30.8% | 26.5% | 100.0% | 100.0% | 80.6% | 53.9% | 1.1% | 100.0% | 3.6% | 46.1% | 4.3% | 100.0% |
| | ZG198 | 19.1% | 16.5% | 100.0% | 100.0% | 63.3% | 98.6% | 4.0% | 100.0% | 7.8% | 100.0% | 30.8% | 38.1% |
| | ZG199 | 100.0% | 6.2% | 36.2% | 100.0% | 63.9% | 46.2% | 5.0% | 100.0% | 5.5% | 24.0% | 3.3% | 100.0% |

Figure S1. **Ion ratios from MRM analysis**. Ion ratios are reported as peak area percentages as compared to the largest transition peak. Values for the heavy isotope standard have been averaged with standard deviation to serve as a model for the light isotope endogenous GVPs. ZG182-184 = E1, ZG185-187 = E2, ZG188-190 = E3, ZG194-196 = A1, ZG197-199 = E2.

| RSID | D1.0017 | D1.0020 | U1.0001 | U1.0003 | U1.0005 |
|---|---|---|---|---|---|
| rs36022742 | C/T | C/T | C/C | C/T | C/C |
| rs1695 | G/G | A/G | A/A | A/G | A/G |
| rs2852464 | C/C | C/C | G/C | G/G | G/G |
| rs1732263 | C/G | C/G | C/C | C/C | C/C |
| rs7212938 | T/T | T/T | T/T | T/T | G/T |
| rs7213256 | C/T | C/C | C/T | C/C | C/C |
| rs17843021 | G/A | A/A | G/G | G/A | G/G |
| rs2071563 | G/A | G/A | G/A | A/A | G/G |
| rs743686 | A/G | G/G | A/G | A/G | A/G |
| rs1455555 | A/G | A/A | A/G | A/G | A/G |
| rs2233391 | C/C | C/C | C/A | A/A | A/A |
| rs10805890 | A/A | A/A | A/A | A/G | A/G |

Table S1. **Genotypes for GVP-associated SNPs from whole exome sequencing**. Each box represents a genotype, and only genotypes of interest are included. D1 series donors are of African ancestry, and U1 series donors are of European ancestry. D1.0017, A1; D1.0020, A2; U1.0001, E1, U1.0003, E2; U1.0005, E3.

| Peptide | Retention Time (min) | Precursor | Transitions | | | | | | | | | | | | | | | | | | | | | | | |
|---|---|---|---|---|---|---|---|---|---|---|---|---|---|---|---|---|---|---|---|---|---|---|---|---|---|---|
| DSQECILTETEAR | 51.1 | 776.3514++ | 1349.6366+ | 1221.5780+ | 1092.5354+ | 932.5047+ | 819.4207+ | 706.3366+ | 605.2889+ | 476.2463+ | 460.1674+ | 620.1981+ | 733.2821+ | 846.3662+ | 947.4139+ | 1076.4565+ | 1177.5041+ | 1306.5467+ | | | | | | | | |
| | | 781.3555++ (heavy) | 1359.6448+ | 1231.5862+ | 1102.5437+ | 942.5130+ | 829.4289+ | 716.3449+ | 615.2972+ | 486.2546+ | 460.1674+ | 620.1981+ | 733.2821+ | 846.3662+ | 947.4139+ | 1076.4565+ | 1177.5041+ | 1306.5467+ | | | | | | | | |
| DSQECILMETEAR | 55.8 | 791.3478++ | 1379.6294+ | 1251.5708+ | 1122.5282+ | 962.4975+ | 849.4135+ | 736.3294+ | 605.2889+ | 476.2463+ | 460.1674+ | 620.1981+ | 733.2821+ | 846.3662+ | 977.4067+ | 1106.4493+ | 1207.4970+ | 1336.5395+ | | | | | | | | |
| | | 796.3519++ (heavy) | 1389.6376+ | 1261.5791+ | 1132.5365+ | 972.5058+ | 859.4217+ | 746.3377+ | 615.2972+ | 486.2546+ | 460.1674+ | 620.1981+ | 733.2821+ | 846.3662+ | 977.4067+ | 1106.4493+ | 1207.4970+ | 1336.5395+ | | | | | | | | |
| GILIDTSR | 45.8 | 437.7533++ | 817.4778+ | 704.3937+ | 591.3097+ | 478.2256+ | 363.1987+ | 284.1969+ | 397.2809+ | 512.3079+ | 613.3556+ | 700.3876+ | | | | | | | | | | | | | | |
| | | 442.7574++ (heavy) | 827.4861+ | 714.4020+ | 601.3179+ | 488.2339+ | 373.2069+ | 284.1969+ | 397.2809+ | 512.3079+ | 613.3556+ | 700.3876+ | | | | | | | | | | | | | | |
| GILVDTSR | 40.9 | 430.7454++ | 803.4621+ | 690.3781+ | 577.2940+ | 478.2256+ | 363.1987+ | 284.1969+ | 383.2653+ | 498.2922+ | 599.3399+ | 686.3719+ | | | | | | | | | | | | | | |
| | | 435.7496++ (heavy) | 813.4704+ | 700.3863+ | 587.3023+ | 488.2339+ | 373.2069+ | 284.1969+ | 383.2653+ | 498.2922+ | 599.3399+ | 686.3719+ | | | | | | | | | | | | | | |
| ALETVQER | 30.4 | 473.2536++ | 874.4629+ | 761.3788+ | 632.3362+ | 531.2885+ | 432.2201+ | 314.1710+ | 415.2187+ | 514.2871+ | 642.3457+ | 771.3883+ | | | | | | | | | | | | | | |
| | | 478.2578++ (heavy) | 884.4711+ | 771.3871+ | 642.3445+ | 541.2968+ | 442.2284+ | 314.1710+ | 415.2187+ | 514.2871+ | 642.3457+ | 771.3883+ | | | | | | | | | | | | | | |
| ALETLQER | 36.5 | 480.2615++ | 888.4785+ | 775.3945+ | 646.3519+ | 545.3042+ | 432.2201+ | 314.1710+ | 415.2187+ | 528.3028+ | 656.3614+ | 785.4040+ | | | | | | | | | | | | | | |
| | | 485.2656++ (heavy) | 898.4868+ | 785.4027+ | 656.3601+ | 555.3125+ | 442.2284+ | 314.1710+ | 415.2187+ | 528.3028+ | 656.3614+ | 785.4040+ | | | | | | | | | | | | | | |
| LEGEINTYR | 40.2 | 547.7775++ | 981.4636+ | 852.4210+ | 795.3995+ | 666.3570+ | 553.2729+ | 439.2300+ | 338.1823+ | 300.1554+ | 429.1980+ | 542.2821+ | 656.3250+ | 757.3727+ | 920.4360+ | | | | | | | | | | | |
| | | 552.7816++ (heavy) | 991.4719+ | 862.4293+ | 805.4078+ | 676.3652+ | 563.2812+ | 449.2382+ | 348.1905+ | 243.1339+ | 300.1554+ | 429.1980+ | 542.2821+ | 656.3250+ | 757.3727+ | 920.4360+ | | | | | | | | | | |
| LEGEINMYR | 47.6 | 562.7739++ | 1011.4564+ | 882.4138+ | 825.3923+ | 696.3498+ | 583.2657+ | 469.2228+ | 338.1823+ | 243.1339+ | 300.1554+ | 429.1980+ | 542.2821+ | 656.3250+ | 787.3655+ | 950.4288+ | | | | | | | | | | |
| | | 567.7780++ (heavy) | 1021.4647+ | 892.4221+ | 835.4006+ | 706.3580+ | 593.2740+ | 479.2310+ | 348.1905+ | 243.1339+ | 300.1554+ | 429.1980+ | 542.2821+ | 656.3250+ | 787.3655+ | 950.4288+ | | | | | | | | | | |
| YISLIYTNYEAGKDDYVK | 59.6 | 1078.0333++ | 1402.6485+ | 1301.6008+ | 1187.5579+ | 1024.4946+ | 895.4520+ | 824.4149+ | 767.3934+ | 639.2984+ | 524.2715+ | 409.2445+ | 364.1867+ | 477.2708+ | 590.3548+ | 753.4182+ | 854.4658+ | 968.5088+ | 1131.5721+ | 1260.6147+ | 1331.6518+ | 1388.6733+ | | | | |
| | | 1082.0404++ (heavy) | 1410.6627+ | 1309.6150+ | 1195.5721+ | 1032.5088+ | 903.4662+ | 832.4291+ | 775.4076+ | 647.3126+ | 532.2857+ | 417.2587+ | 364.1867+ | 477.2708+ | 590.3548+ | 753.4182+ | 854.4658+ | 968.5088+ | 1131.5721+ | 1260.6147+ | 1331.6518+ | 1388.6733+ | | | | |
| YVSLIYTNYEAGKDDYVK | 57.7 | 1071.0255++ | 1402.6485+ | 1301.6008+ | 1187.5579+ | 1024.4946+ | 895.4520+ | 824.4149+ | 767.3934+ | 639.2984+ | 524.2715+ | 409.2445+ | 350.1710+ | 463.2551+ | 576.3392+ | 739.4025+ | 840.4502+ | 954.4931+ | 1117.5564+ | 1246.5990+ | 1317.6361+ | 1374.6576+ | | | | |
| | | 1075.0326++ (heavy) | 1410.6627+ | 1309.6150+ | 1195.5721+ | 1032.5088+ | 903.4662+ | 832.4291+ | 775.4076+ | 647.3126+ | 532.2857+ | 417.2587+ | 350.1710+ | 463.2551+ | 576.3392+ | 739.4025+ | 840.4502+ | 954.4931+ | 1117.5564+ | 1246.5990+ | 1317.6361+ | 1374.6576+ | | | | |
| VSAMYSSSSCKLPSLSPVAR | 53.8 | 709.6906+++ | 1488.7839+ | 1401.7519+ | 1314.7198+ | 1227.6878+ | 1067.6572+ | 939.5622+ | 826.4781+ | 729.4254+ | 642.3933+ | 529.3093+ | 442.2772+ | 345.2245+ | 258.1448+ | 389.1853+ | 552.2486+ | 639.2807+ | 726.3127+ | 813.3447+ | 900.3768+ | 1060.4074+ | 1188.5024+ | 1301.5864+ | 1398.6392+ | 1485.6712+ |
| | | 713.0267+++ (heavy) | 1498.7922+ | 1411.7601+ | 1324.7281+ | 1237.6961+ | 1077.6654+ | 949.5705+ | 836.4864+ | 739.4336+ | 652.4016+ | 539.3175+ | 452.2855+ | 355.2327+ | 258.1448+ | 389.1853+ | 552.2486+ | 639.2807+ | 726.3127+ | 813.3447+ | 900.3768+ | 1060.4074+ | 1188.5024+ | 1301.5864+ | 1398.6392+ | 1485.6712+ |
| VSAMYSSSPCKLPSLSPVAR | 55.0 | 713.0309+++ | 1411.7726+ | 1324.7406+ | 1227.6878+ | 1067.6572+ | 939.5622+ | 826.4781+ | 729.4254+ | 642.3933+ | 529.3093+ | 442.2772+ | 345.2245+ | 258.1448+ | 389.1853+ | 552.2486+ | 639.2807+ | 726.3127+ | 813.3447+ | 910.3975+ | 1070.4281+ | 1198.5231+ | 1311.6072+ | 1408.6599+ | 1495.6920+ | |
| | | 716.3669+++ (heavy) | 1421.7809+ | 1334.7488+ | 1237.6961+ | 1077.6654+ | 949.5705+ | 836.4864+ | 739.4336+ | 652.4016+ | 539.3175+ | 452.2855+ | 355.2327+ | 258.1448+ | 389.1853+ | 552.2486+ | 639.2807+ | 726.3127+ | 813.3447+ | 910.3975+ | 1070.4281+ | 1198.5231+ | 1311.6072+ | 1408.6599+ | 1495.6920+ | |
| EHCSACGPLSR | 27.8 | 637.2742++ | 1144.4986+ | 1007.4397+ | 847.4091+ | 760.3770+ | 689.3399+ | 529.3093+ | 472.2878+ | 375.2350+ | 427.1394+ | 514.1715+ | 585.2086+ | 745.2392+ | 802.2607+ | 899.3134+ | 1012.3975+ | 1099.4295+ | | | | | | | | |
| | | 642.2784++ (heavy) | 1154.5069+ | 1017.4480+ | 857.4173+ | 770.3853+ | 699.3482+ | 539.3175+ | 482.2961+ | 385.2433+ | 427.1394+ | 514.1715+ | 585.2086+ | 745.2392+ | 802.2607+ | 899.3134+ | 1012.3975+ | 1099.4295+ | | | | | | | | |
| EHCSACGPLSQLLVK | 55.6 | 849.9187++ | 1432.7287+ | 1272.6980+ | 1185.6660+ | 1114.6289+ | 954.5982+ | 897.5768+ | 800.5240+ | 687.4400+ | 600.4079+ | 472.3493+ | 359.2653+ | 427.1394+ | 514.1715+ | 585.2086+ | 745.2392+ | 802.2607+ | 899.3134+ | 1012.3975+ | 1099.4295+ | 1227.4881+ | 1340.5722+ | 1453.6562+ | | |
| | | 853.9258++ (heavy) | 1440.7429+ | 1280.7122+ | 1193.6802+ | 1122.6431+ | 962.6124+ | 905.5910+ | 808.5382+ | 695.4542+ | 608.4221+ | 480.3635+ | 367.2795+ | 427.1394+ | 514.1715+ | 585.2086+ | 745.2392+ | 802.2607+ | 899.3134+ | 1012.3975+ | 1099.4295+ | 1227.4881+ | 1340.5722+ | 1453.6562+ | | |
| EWSTFAVGPGHCLQLNDR | 60.3 | 1043.9918++ | 1436.7063+ | 1365.6692+ | 1266.6008+ | 1209.5793+ | 1112.5266+ | 1055.5051+ | 918.4462+ | 758.4155+ | 645.3315+ | 517.2729+ | 404.1888+ | 403.1612+ | 504.2089+ | 651.2773+ | 722.3144+ | 821.3828+ | 878.4043+ | 975.4571+ | 1032.4785+ | 1169.5374+ | 1329.5681+ | 1442.6521+ | | |
| | | 1048.9959++ (heavy) | 1446.7146+ | 1375.6775+ | 1276.6090+ | 1219.5876+ | 1122.5348+ | 1065.5134+ | 928.4544+ | 768.4238+ | 655.3397+ | 527.2812+ | 414.1971+ | 403.1612+ | 504.2089+ | 651.2773+ | 722.3144+ | 821.3828+ | 878.4043+ | 975.4571+ | 1032.4785+ | 1169.5374+ | 1329.5681+ | 1442.6521+ | | |
| EWSTFAVGPGHCLQLHDR | 56.0 | 1055.4998++ | 1459.7223+ | 1388.6852+ | 1289.6168+ | 1232.5953+ | 1135.5425+ | 1078.5211+ | 941.4622+ | 781.4315+ | 668.3474+ | 540.2889+ | 427.2048+ | 403.1612+ | 504.2089+ | 651.2773+ | 722.3144+ | 821.3828+ | 878.4043+ | 975.4571+ | 1032.4785+ | 1169.5374+ | 1329.5681+ | 1442.6521+ | | |
| | | 1060.5039++ (heavy) | 1469.7306+ | 1398.6934+ | 1299.6250+ | 1242.6036+ | 1145.5508+ | 1088.5293+ | 951.4704+ | 791.4398+ | 678.3557+ | 550.2971+ | 437.2131+ | 403.1612+ | 504.2089+ | 651.2773+ | 722.3144+ | 821.3828+ | 878.4043+ | 975.4571+ | 1032.4785+ | 1169.5374+ | 1329.5681+ | 1442.6521+ | | |
| GVALSNVIHK | 41.3 | 519.3087++ | 980.5887+ | 881.5203+ | 810.4832+ | 697.3991+ | 610.3671+ | 496.3242+ | 397.2558+ | 228.1343+ | 341.2183+ | 428.2504+ | 542.2933+ | 641.3617+ | 754.4458+ | 891.5047+ | | | | | | | | | | |
| | | 523.3158++ (heavy) | 988.6029+ | 889.5345+ | 818.4974+ | 705.4133+ | 618.3813+ | 504.3384+ | 405.2700+ | 228.1343+ | 341.2183+ | 428.2504+ | 542.2933+ | 641.3617+ | 754.4458+ | 891.5047+ | | | | | | | | | | |
| GVALSNVVHK | 36.6 | 512.3009++ | 966.5731+ | 867.5047+ | 796.4676+ | 683.3835+ | 596.3515+ | 482.3085+ | 383.2401+ | 228.1343+ | 341.2183+ | 428.2504+ | 542.2933+ | 641.3617+ | 740.4301+ | 877.4890+ | | | | | | | | | | |
| | | 516.3080++ (heavy) | 974.5873+ | 875.5189+ | 804.4818+ | 691.3977+ | 604.3657+ | 490.3227+ | 391.2543+ | 228.1343+ | 341.2183+ | 428.2504+ | 542.2933+ | 641.3617+ | 740.4301+ | 877.4890+ | | | | | | | | | | |
| DLNMDCMVAEIK | 64.9 | 719.8224++ | 1323.6105+ | 1210.5265+ | 1096.4835+ | 965.4431+ | 850.4161+ | 690.3855+ | 559.3450+ | 460.2766+ | 389.2395+ | 343.1612+ | 474.2017+ | 589.2286+ | 749.2593+ | 880.2998+ | 979.3682+ | 1050.4053+ | 1179.4479+ | 1292.5320+ | | | | | | |
| | | 723.8295++ (heavy) | 1331.6247+ | 1218.5407+ | 1104.4977+ | 973.4573+ | 858.4303+ | 698.3997+ | 567.3592+ | 468.2908+ | 397.2537+ | 343.1612+ | 474.2017+ | 589.2286+ | 749.2593+ | 880.2998+ | 979.3682+ | 1050.4053+ | 1179.4479+ | 1292.5320+ | | | | | | |
| DLNMDCIVAEIK | 67.3 | 710.8442++ | 1305.6541+ | 1192.5701+ | 1078.5271+ | 947.4866+ | 832.4597+ | 672.4291+ | 559.3450+ | 460.2766+ | 389.2395+ | 343.1612+ | 474.2017+ | 589.2286+ | 749.2593+ | 862.3434+ | 961.4118+ | 1032.4489+ | 1161.4915+ | 1274.5755+ | | | | | | |
| | | 714.8513++ (heavy) | 1313.6683+ | 1200.5843+ | 1086.5413+ | 955.5008+ | 840.4739+ | 680.4433+ | 567.3592+ | 468.2908+ | 397.2537+ | 343.1612+ | 474.2017+ | 589.2286+ | 749.2593+ | 862.3434+ | 961.4118+ | 1032.4489+ | 1161.4915+ | 1274.5755+ | | | | | | |
| GAFLYEPCGVSTPVLSTGVLR | 67.1 | 1112.0775++ | 1445.7781+ | 1285.7474+ | 1228.7260+ | 1129.6575+ | 1042.6255+ | 941.5778+ | 844.5251+ | 745.4567+ | 632.3726+ | 545.3406+ | 444.2929+ | 387.2714+ | 276.1343+ | 389.2183+ | 552.2817+ | 681.3243+ | 778.3770+ | 938.4077+ | 995.4291+ | 1094.4975+ | 1181.5296+ | 1282.5773+ | 1379.6300+ | 1478.6984+ |
| | | 1117.0817++ (heavy) | 1455.7863+ | 1295.7557+ | 1238.7342+ | 1139.6658+ | 1052.6338+ | 951.5861+ | 854.5333+ | 755.4649+ | 642.3809+ | 555.3488+ | 454.3012+ | 397.2797+ | 276.1343+ | 389.2183+ | 552.2817+ | 681.3243+ | 778.3770+ | 938.4077+ | 995.4291+ | 1094.4975+ | 1181.5296+ | 1282.5773+ | 1379.6300+ | 1478.6984+ |
| GAFLYDPCGVSTPVLSTGVLR | 67.7 | 1105.0697++ | 1445.7781+ | 1285.7474+ | 1228.7260+ | 1129.6575+ | 1042.6255+ | 941.5778+ | 844.5251+ | 745.4567+ | 632.3726+ | 545.3406+ | 444.2929+ | 387.2714+ | 276.1343+ | 389.2183+ | 552.2817+ | 667.3086+ | 764.3614+ | 924.3920+ | 981.4135+ | 1080.4819+ | 1167.5139+ | 1268.5616+ | 1365.6144+ | 1464.6828+ |
| | | 1110.0739++ (heavy) | 1455.7863+ | 1295.7557+ | 1238.7342+ | 1139.6658+ | 1052.6338+ | 951.5861+ | 854.5333+ | 755.4649+ | 642.3809+ | 555.3488+ | 454.3012+ | 397.2797+ | 276.1343+ | 389.2183+ | 552.2817+ | 667.3086+ | 764.3614+ | 924.3920+ | 981.4135+ | 1080.4819+ | 1167.5139+ | 1268.5616+ | 1365.6144+ | 1464.6828+ |
| ARPLEQAVAAIVCTFQEYAGR | 75.6 | 1196.6075++ | 1485.7155+ | 1414.6784+ | 1343.6412+ | 1230.5572+ | 1131.4888+ | 971.4581+ | 870.4104+ | 723.3420+ | 595.2835+ | 466.2409+ | 303.1775+ | 367.2088+ | 480.2929+ | 609.3355+ | 737.3941+ | 808.4312+ | 907.4996+ | 978.5367+ | 1049.5738+ | 1162.6579+ | 1261.7263+ | 1421.7569+ | | |
| | | 1201.6117++ (heavy) | 1495.7237+ | 1424.6866+ | 1353.6495+ | 1240.5655+ | 1141.4970+ | 981.4664+ | 880.4187+ | 733.3503+ | 605.2917+ | 476.2491+ | 313.1858+ | 367.2088+ | 480.2929+ | 609.3355+ | 737.3941+ | 808.4312+ | 907.4996+ | 978.5367+ | 1049.5738+ | 1162.6579+ | 1261.7263+ | 1421.7569+ | | |
| AKPLEQAVAAIVCTFQEYAGR | N/A | 1182.6045++ | 1485.7155+ | 1414.6784+ | 1343.6412+ | 1230.5572+ | 1131.4888+ | 971.4581+ | 870.4104+ | 723.3420+ | 595.2835+ | 466.2409+ | 303.1775+ | 339.2027+ | 452.2867+ | 581.3293+ | 709.3879+ | 780.4250+ | 879.4934+ | 950.5306+ | 1021.5677+ | 1134.6517+ | 1233.7201+ | 1393.7508+ | 1494.7985+ | |
| | | 1187.6086++ (heavy) | 1495.7237+ | 1424.6866+ | 1353.6495+ | 1240.5655+ | 1141.4970+ | 981.4664+ | 880.4187+ | 733.3503+ | 605.2917+ | 476.2491+ | 313.1858+ | 339.2027+ | 452.2867+ | 581.3293+ | 709.3879+ | 780.4250+ | 879.4934+ | 950.5306+ | 1021.5677+ | 1134.6517+ | 1233.7201+ | 1393.7508+ | 1494.7985+ | |

Table S2. **Transitions list for PRM analysis**. A list of transitions that were used to identify endogenous and isotopically-labeled GVPs. Transitions were chosen based on appearance in standard peptide spectra.

| Peptide | Retention Time (min) | Precursor | Transitions | | |
|---|---|---|---|---|---|
| DSQECILTETEAR | 4.7 | 776.3514++ | 819.4207+ | 706.3366+ | 460.1674+ |
| | | 781.3555++ (heavy) | 829.4289+ | 716.3449+ | 460.1674+ |
| DSQECILMETEAR | 5.4 | 791.3478++ | 849.4135+ | 736.3294+ | 460.1674+ |
| | | 796.3519++ (heavy) | 859.4217+ | 746.3377+ | 460.1674+ |
| GILIDTSR | 4.8 | 437.7533++ | 704.3937+ | 591.3097+ | 284.1969+ |
| | | 442.7574++ (heavy) | 714.4020+ | 601.3179+ | 284.1969+ |
| GILVDTSR | 4.1 | 430.7454++ | 577.2940+ | 478.2256+ | 284.1969+ |
| | | 435.7496++ (heavy) | 587.3023+ | 488.2339+ | 284.1969+ |
| ALETVQER | 2.5 | 473.2536++ | 632.3362+ | 531.2885+ | 314.1710+ |
| | | 478.2578++ (heavy) | 642.3445+ | 541.2968+ | 314.1710+ |
| ALETLQER | 3.4 | 480.2615++ | 646.3519+ | 545.3042+ | 314.1710+ |
| | | 485.2656++ (heavy) | 656.3601+ | 555.3125+ | 314.1710+ |
| LEGEINTYR | 3.6 | 547.7775++ | 981.4636+ | 852.4210+ | 243.1339+ |
| | | 552.7816++ (heavy) | 991.4719+ | 862.4293+ | 243.1339+ |
| LEGEINMYR | 4.7 | 562.7739++ | 1011.4564+ | 882.4138+ | 243.1339+ |
| | | 567.7780++ (heavy) | 1021.4647+ | 892.4221+ | 243.1339+ |
| YISLIYTNYEAGKDDYVK | 6.9 | 719.0246+++ | 895.4520+ | 277.1547+ | 364.1867+ |
| | | 721.6960+++ (heavy) | 903.4662+ | 277.1547+ | 364.1867+ |
| YVSLIYTNYEAGKDDYVK | 6.4 | 1071.0255++ | 1402.6485+ | 263.1390+ | 350.1710+ |
| | | 1075.0326++ (heavy) | 1410.6627+ | 263.1390+ | 350.1710+ |
| VSAMYSSSSCKLPSLSPVAR | 6.0 | 709.6906+++ | 826.4781+ | 529.3093+ | 258.1448+ |
| | | 713.0267+++ (heavy) | 836.4864+ | 539.3175+ | 258.1448+ |
| VSAMYSSSPCKLPSLSPVAR | 6.2 | 713.0309+++ | 826.4781+ | 529.3093+ | 258.1448+ |
| | | 716.3669+++ (heavy) | 836.4864+ | 539.3175+ | 258.1448+ |
| EHCSACGPLSR | 2.1 | 637.2742++ | 1007.4397+ | 847.4091+ | 427.1394+ |
| | | 642.2784++ (heavy) | 1017.4480+ | 857.4173+ | 427.1394+ |
| EHCSACGPLSQLLVK | 6.8 | 849.9187++ | 1432.7287+ | 954.5982+ | 427.1394+ |
| | | 853.9258++ (heavy) | 1440.7429+ | 962.6124+ | 427.1394+ |
| EWSTFAVGPGHCLQLNDR | 7.0 | 696.3303+++ | 645.3315+ | 517.2729+ | 722.3144+ |
| | | 699.6664+++ (heavy) | 655.3397+ | 527.2812+ | 722.3144+ |
| EWSTFAVGPGHCLQLHDR | 6.3 | 704.0023+++ | 1289.6168+ | 427.2048+ | 722.3144+ |
| | | 707.3384+++ (heavy) | 1299.6250+ | 437.2131+ | 722.3144+ |
| GVALSNVIHK | 4.6 | 519.3087++ | 697.3991+ | 397.2558+ | 228.1343+ |
| | | 523.3158++ (heavy) | 705.4133+ | 405.2700+ | 228.1343+ |
| GVALSNVVHK | 3.9 | 512.3009++ | 867.5047+ | 796.4676+ | 683.3835+ |
| | | 516.3080++ (heavy) | 875.5189+ | 804.4818+ | 691.3977+ |
| DLNMDCMVAEIK | 7.6 | 719.8224++ | 965.4431+ | 690.3855+ | 343.1612+ |
| | | 723.8295++ (heavy) | 973.4573+ | 698.3997+ | 343.1612+ |
| DLNMDCIVAEIK | 8.4 | 710.8442++ | 947.4866+ | 672.4291+ | 343.1612+ |
| | | 714.8513++ (heavy) | 955.5008+ | 680.4433+ | 343.1612+ |
| GAFLYEPCGVSTPVLSTGVLR | 8.9 | 1112.0775++ | 941.5778+ | 681.3243+ | 1094.4975+ |
| | | 1117.0817++ (heavy) | 951.5861+ | 681.3243+ | 1094.4975+ |
| GAFLYDPCGVSTPVLSTGVLR | 9.0 | 1105.0697++ | 941.5778+ | 667.3086+ | 1080.4819+ |
| | | 1110.0739++ (heavy) | 951.5861+ | 667.3086+ | 1080.4819+ |
| ARPLEQAVAAIVCTFQEYAGR | 12.5 | 798.0741+++ | 1485.7155+ | 270.1561+ | 1162.6579+ |
| | | 801.4102+++ (heavy) | 1495.7237+ | 270.1561+ | 1162.6579+ |
| AKPLEQAVAAIVCTFQEYAGR | 12.5 | 1182.6045++ | 1485.7155+ | 242.1499+ | 1134.6517+ |
| | | 1187.6086++ (heavy) | 1495.7237+ | 242.1499+ | 1134.6517+ |

Table S3. **Transitions list for MRM analysis**. A list of transitions that were used to identify endogenous and isotopically-labeled GVPs. Transitions were chosen based on appearance in standard peptide spectra.

| RAW file name | Sample ID | Donor | Background | Analytical Method |
|---|---|---|---|---|
| QEPlus2_03162018_34_ZG182.raw | ZG182 | U1.0001 | European | QE+ |
| QEPlus2_03162018_84_ZG183.raw | ZG183 | U1.0001 | European | QE+ |
| QEPlus2_03162018_93_ZG184.raw | ZG184 | U1.0001 | European | QE+ |
| QEPlus2_03162018_24_ZG185.raw | ZG185 | U1.0003 | European | QE+ |
| QEPlus2_03162018_32_ZG186.raw | ZG186 | U1.0003 | European | QE+ |
| QEPlus2_03162018_28_ZG187.raw | ZG187 | U1.0003 | European | QE+ |
| QEPlus2_03162018_39_ZG188.raw | ZG188 | U1.0005 | European | QE+ |
| QEPlus2_03162018_89_ZG189_180326120408.raw | ZG189 | U1.0005 | European | QE+ |
| QEPlus2_03162018_59_ZG190.raw | ZG190 | U1.0005 | European | QE+ |
| QEPlus2_03162018_61_ZG194.raw | ZG194 | D1.0017 | African | QE+ |
| QEPlus2_03162018_68_ZG195.raw | ZG195 | D1.0017 | African | QE+ |
| QEPlus2_03162018_22_ZG196.raw | ZG196 | D1.0017 | African | QE+ |
| QEPlus2_03162018_78_ZG197.raw | ZG197 | D1.0020 | African | QE+ |
| QEPlus2_03162018_72_ZG198.raw | ZG198 | D1.0020 | African | QE+ |
| QEPlus2_03162018_70_ZG199.raw | ZG199 | D1.0020 | African | QE+ |
| QEPlus2_03162018_18_ZG200.raw | ZG200 | Blank with Trypsin | N/A | QE+ |
| QEPlus2_03162018_20_ZG201.raw | ZG201 | Blank without Trypsin | N/A | QE+ |
| FL05222020_28_ZG182.raw | ZG182 | U1.0001 | European | Lumos (DIA) |
| FL05222020_36_ZG183.raw | ZG183 | U1.0001 | European | Lumos (DIA) |
| FL05222020_50_ZG184.raw | ZG184 | U1.0001 | European | Lumos (DIA) |
| FL05222020_58_ZG185.raw | ZG185 | U1.0003 | European | Lumos (DIA) |
| FL05222020_44_ZG186.raw | ZG186 | U1.0003 | European | Lumos (DIA) |
| FL05222020_48_ZG187.raw | ZG187 | U1.0003 | European | Lumos (DIA) |
| FL05222020_60_ZG188.raw | ZG188 | U1.0005 | European | Lumos (DIA) |
| FL05222020_34_ZG189.raw | ZG189 | U1.0005 | European | Lumos (DIA) |
| FL05222020_54_ZG190.raw | ZG190 | U1.0005 | European | Lumos (DIA) |
| FL05222020_30_ZG194.raw | ZG194 | D1.0017 | African | Lumos (DIA) |
| FL05222020_40_ZG195.raw | ZG195 | D1.0017 | African | Lumos (DIA) |
| FL05222020_62_ZG196.raw | ZG196 | D1.0017 | African | Lumos (DIA) |
| FL05222020_66_ZG197.raw | ZG197 | D1.0020 | African | Lumos (DIA) |
| FL05222020_42_ZG198.raw | ZG198 | D1.0020 | African | Lumos (DIA) |
| FL05222020_64_ZG199.raw | ZG199 | D1.0020 | African | Lumos (DIA) |
| FL05222020_52_ZG200.raw | ZG200 | Blank with Trypsin | N/A | Lumos (DIA) |
| FL05222020_32_ZG201.raw | ZG201 | Blank without Trypsin | N/A | Lumos (DIA) |
| FL20180724_40_ZG182.raw | ZG182 | U1.0001 | European | Lumos (QD) |
| FL20180724_44_ZG183.raw | ZG183 | U1.0001 | European | Lumos (QD) |
| FL20180724_42_ZG184.raw | ZG184 | U1.0001 | European | Lumos (QD) |
| FL20180724_32_ZG185.raw | ZG185 | U1.0003 | European | Lumos (QD) |
| FL20180724_22_ZG186.raw | ZG186 | U1.0003 | European | Lumos (QD) |
| FL20180724_11_ZG187.raw | ZG187 | U1.0003 | European | Lumos (QD) |
| FL20180724_14_zg188.raw | ZG188 | U1.0005 | European | Lumos (QD) |
| FL20180724_30_ZG189.raw | ZG189 | U1.0005 | European | Lumos (QD) |
| FL20180724_26_ZG190.raw | ZG190 | U1.0005 | European | Lumos (QD) |
| FL20180724_36_ZG194.raw | ZG194 | D1.0017 | African | Lumos (QD) |
| FL20180724_20_ZG195.raw | ZG195 | D1.0017 | African | Lumos (QD) |
| FL20180724_48_ZG196.raw | ZG196 | D1.0017 | African | Lumos (QD) |
| FL20180724_16_ZG197.raw | ZG197 | D1.0020 | African | Lumos (QD) |
| FL20180724_55_ZG198REAL.raw | ZG198 | D1.0020 | African | Lumos (QD) |
| FL20180724_38_ZG199.raw | ZG199 | D1.0020 | African | Lumos (QD) |
| FL20180724_57_ZG200REAL.raw | ZG200 | Blank with Trypsin | N/A | Lumos (QD) |
| FL20180724_34_ZG201.raw | ZG201 | Blank without Trypsin | N/A | Lumos (QD) |
| ZG182.d | ZG182 | U1.0001 | European | QQQ |
| ZG183.d | ZG183 | U1.0001 | European | QQQ |
| ZG184.d | ZG184 | U1.0001 | European | QQQ |
| ZG185.d | ZG185 | U1.0003 | European | QQQ |
| ZG186.d | ZG186 | U1.0003 | European | QQQ |
| ZG187.d | ZG187 | U1.0003 | European | QQQ |
| ZG188.d | ZG188 | U1.0005 | European | QQQ |
| ZG189.d | ZG189 | U1.0005 | European | QQQ |
| ZG190.d | ZG190 | U1.0005 | European | QQQ |
| ZG194.d | ZG194 | D1.0017 | African | QQQ |
| ZG195.d | ZG195 | D1.0017 | African | QQQ |
| ZG196.d | ZG196 | D1.0017 | African | QQQ |
| ZG197.d | ZG197 | D1.0020 | African | QQQ |
| ZG198.d | ZG198 | D1.0020 | African | QQQ |
| ZG199.d | ZG199 | D1.0020 | African | QQQ |
| ZG200.d | ZG200 | Blank with Trypsin | N/A | QQQ |
| ZG201.d | ZG201 | Blank without Trypsin | N/A | QQQ |

Table S4. **Sample master list**. This table includes all samples that were mentioned in this research. All samples used for this manuscript were compiled and uploaded to proteome exchange (PDX_______) and are available online.

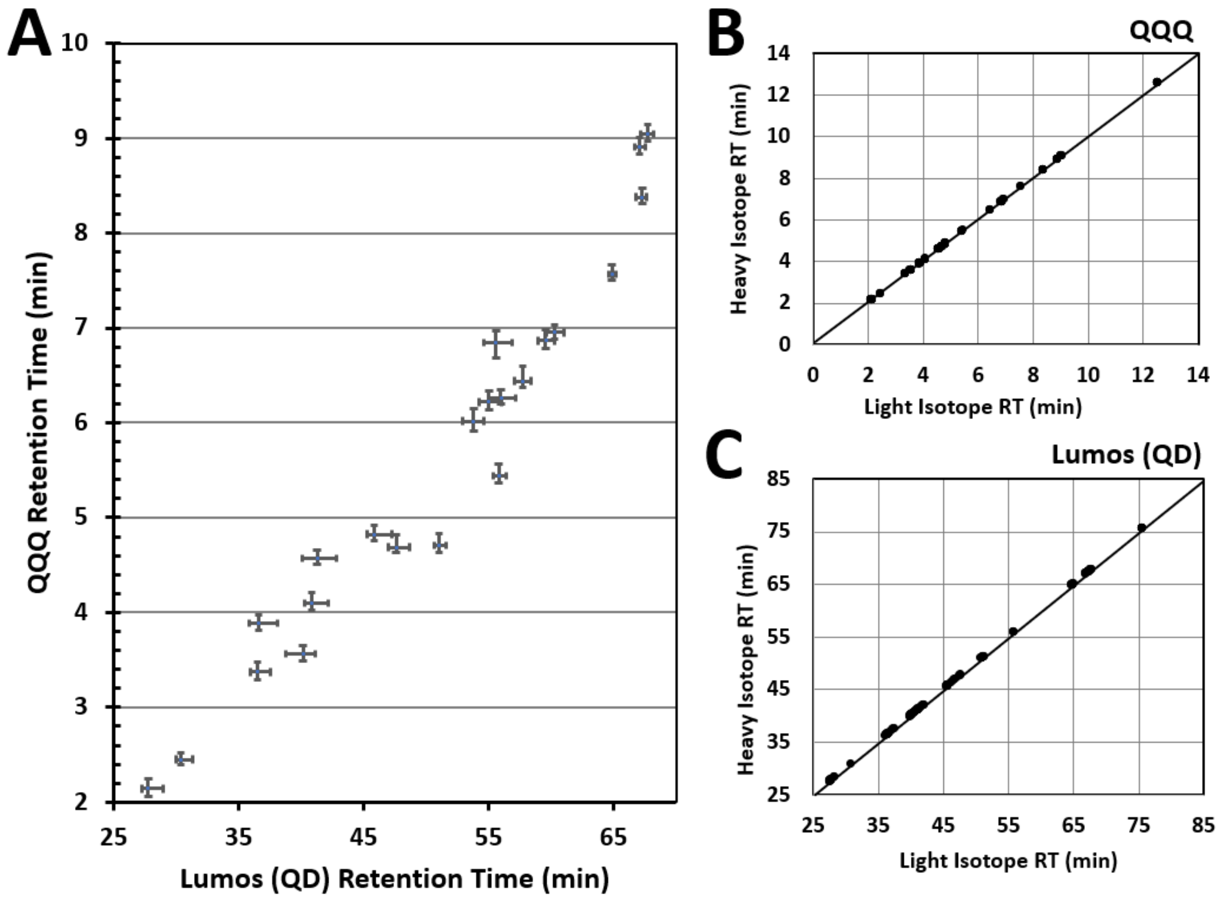


Figure S2. **Retention time comparison for targeted proteomic methods**. Tighter error bars and more resolved data points represents less significant retention time shifting. **A**) Comparing retention times for MRM and PRM. Peptide pair ARPLEQAVAAIVCTFQEYAGR and AKPLEQAVAAIVCTFQEYAGR were left out because one of them is not found in the PRM. Error bars represent variance of apex retention times among 15 samples. **B**) Comparing heavy and light isotope GVP retention time during MRM analysis. **C**) Comparing heavy and light isotope GVP retention time during PRM analysis.

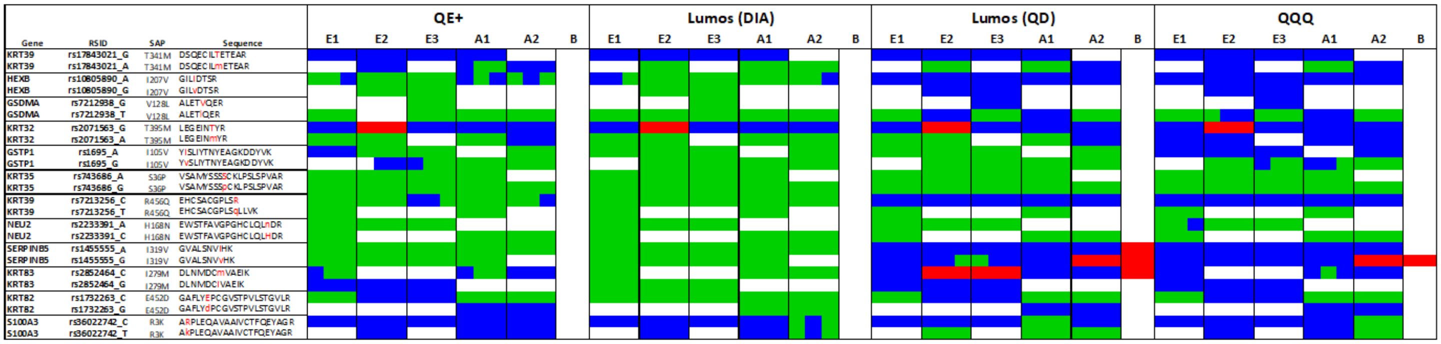


Figure S3. **Expanded GVP matrix comparing four analytical methods of acquisition**. This matrix represents GVPs that have been verified via whole exome sequencing. Each row is a variant peptide and each column is a GVP profile for each of the three replicates. DDA, data dependent acquisition on Q Exactive+; DIA, data independent acquisition on Fusion Lumos; PRM, parallel reaction monitoring on Fusion Lumos; MRM, multiple reaction monitoring on Agilent 6495; E1-3, three European subjects; A1-2, two African subjects.

**A**

| Standard Peptide Sequence | Final amount (nmol) |
|---|---|
| DSQECILTETEAR | 0.25 |
| DSQECILMETEAR | 1.20 |
| GILIDTSR | 1.88 |
| GILVDTSR | 0.79 |
| ALETVQER | 0.23 |
| ALETLQER | 0.53 |
| LEGEINTYR | 0.24 |
| LEGEINMYR | 3.72 |
| YISLIYTNYEAGKDDYVK | 9.75 |
| YVSLIYTNYEAGKDDYVK | 4.71 |
| VSAMYSSSSCKLPSLSPVAR | 4.71 |
| VSAMYSSSPCKLPSLSPVAR | 9.75 |
| EHCSACGPLSR | 1.15 |
| EHCSACGPLSQLLVK | 2.26 |
| EWSTFAVGPGHCLQLNDR | 9.75 |
| EWSTFAVGPGHCLQLHDR | 9.75 |
| GVALSNVIHK | 1.10 |
| GVALSNVVHK | 1.31 |
| DLNMDCMVAEIK | 0.44 |
| DLNMDCIVAEIK | 0.27 |
| GAFLYEPCGVSTPVLSTGVLR | 0.31 |
| GAFLYDPCGVSTPVLSTGVLR | 0.22 |
| A*RPLEQAVAAIVCTFQEYAGR | 6.03 |
| A*KPLEQAVAAIVCTFQEYAGR | 9.75 |

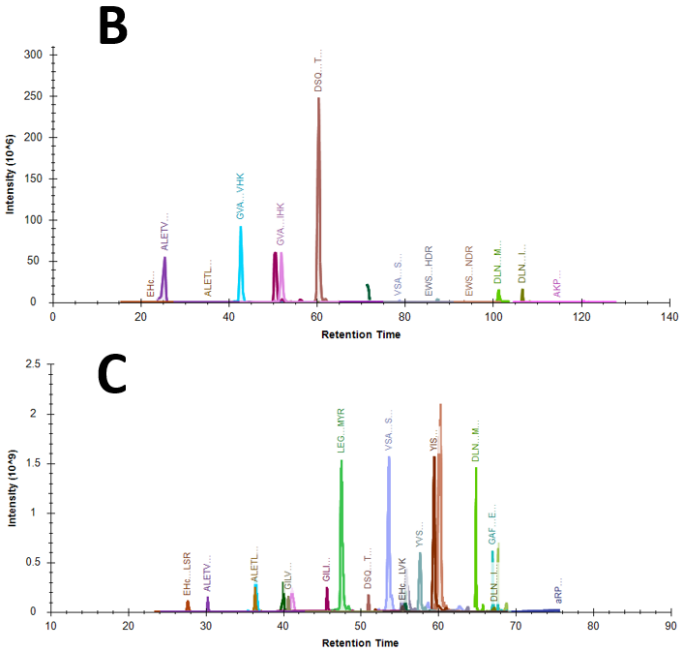


Figure S4. **Normalized peptide standard mixture**. Peptides were normalized to the average peak area among all 24 peptides based on peak areas from a spike mixture of uniform peptide concentrations. **A**) Final peptide amounts in the normalized spike mixture. **B**) Total ion chromatogram of the uniform pre-normalized spike mixture exhibiting high and low peak areas. **C**) Total ion chromatogram of the normalized spike mixture exhibiting more comparable peak areas.

| | DSQECILTETEAR | | | GILIDTSR | | | ALETVQER | | | LEGEINTYR | | |
|---|---|---|---|---|---|---|---|---|---|---|---|---|
| Replicate | y7 - 819.4207+ | y6 - 706.3366+ | b4 - 460.1674+ | y6 - 704.3937+ | y5 - 591.3097+ | b3 - 284.1969+ | y5 - 632.3362+ | y4 - 531.2885+ | b3 - 314.1710+ | y8 - 981.4636+ | y7 - 852.4210+ | b2 - 243.1339+ |
| ZG182 | >>3:1 | >>3:1 | >>3:1 | >>3:1 | >>3:1 | >>3:1 | <<3:1 | <<3:1 | <<3:1 | >>3:1 | >>3:1 | >>3:1 |
| ZG183 | >>3:1 | >>3:1 | >>3:1 | >>3:1 | >>3:1 | >>3:1 | <<3:1 | <<3:1 | <<3:1 | >>3:1 | >>3:1 | >>3:1 |
| ZG184 | >>3:1 | >>3:1 | >>3:1 | >>3:1 | >>3:1 | >>3:1 | <<3:1 | <<3:1 | <<3:1 | >>3:1 | >>3:1 | >>3:1 |
| ZG185 | >>3:1 | >>3:1 | >>3:1 | 13.2:1 | 8.1:1 | 3.4:1 | <<3:1 | <<3:1 | <<3:1 | >>3:1 | >>3:1 | >>3:1 |
| ZG186 | >>3:1 | >>3:1 | >>3:1 | 20.2:1 | 5.7:1 | 5.5:1 | <<3:1 | <<3:1 | <<3:1 | >>3:1 | >>3:1 | >>3:1 |
| ZG187 | >>3:1 | >>3:1 | >>3:1 | 14.9:1 | 11.4:1 | 7.6:1 | <<3:1 | <<3:1 | <<3:1 | >>3:1 | >>3:1 | >>3:1 |
| ZG188 | >>3:1 | >>3:1 | >>3:1 | 14.3:1 | 15.7:1 | 10.7:1 | 5.2:1 | 2.9:1 | 2.1:1 | >>3:1 | >>3:1 | >>3:1 |
| ZG189 | >>3:1 | >>3:1 | >>3:1 | 15.1:1 | 16.8:1 | 5.7:1 | 4.7:1 | 2.2:1 | 2.4:1 | >>3:1 | >>3:1 | >>3:1 |
| ZG190 | >>3:1 | >>3:1 | >>3:1 | 12.7:1 | 9.8:1 | 5.2:1 | 5.2:1 | 0.9:1 | 2.7:1 | >>3:1 | >>3:1 | >>3:1 |
| ZG194 | >>3:1 | >>3:1 | >>3:1 | 11.2:1 | 20.4:1 | 5.4:1 | <<3:1 | <<3:1 | <<3:1 | >>3:1 | >>3:1 | >>3:1 |
| ZG195 | >>3:1 | >>3:1 | >>3:1 | 3.2:1 | 5.8:1 | 4.1:1 | <<3:1 | <<3:1 | <<3:1 | >>3:1 | >>3:1 | >>3:1 |
| ZG196 | >>3:1 | >>3:1 | >>3:1 | 10.4:1 | 11.6:1 | 6.0:1 | <<3:1 | <<3:1 | <<3:1 | >>3:1 | >>3:1 | >>3:1 |
| ZG197 | <<3:1 | <<3:1 | <<3:1 | 6.8:1 | 6.2:1 | 4.3:1 | <<3:1 | <<3:1 | <<3:1 | >>3:1 | >>3:1 | >>3:1 |
| ZG198 | <<3:1 | <<3:1 | <<3:1 | 3.0:1 | 2.0:1 | 4.3:1 | <<3:1 | <<3:1 | <<3:1 | >>3:1 | >>3:1 | >>3:1 |
| ZG199 | <<3:1 | <<3:1 | <<3:1 | 18.7:1 | 8.0:1 | 9.0:1 | <<3:1 | <<3:1 | <<3:1 | >>3:1 | >>3:1 | >>3:1 |

| | DSQECILMETEAR | | | GILVDTSR | | | ALETLQER | | | LEGEINMYR | | |
|---|---|---|---|---|---|---|---|---|---|---|---|---|
| Replicate | y7 - 849.4135+ | y6 - 736.3294+ | b4 - 460.1674+ | y5 - 577.2940+ | y4 - 478.2256+ | b3 - 284.1969+ | y5 - 646.3519+ | y4 - 545.3042+ | b3 - 314.1710+ | y8 - 1011.4564+ | y7 - 882.4138+ | b2 - 243.1339+ |
| ZG182 | <<3:1 | <<3:1 | <<3:1 | <<3:1 | <<3:1 | <<3:1 | <<3:1 | <<3:1 | <<3:1 | >>3:1 | >>3:1 | >>3:1 |
| ZG183 | <<3:1 | <<3:1 | <<3:1 | <<3:1 | <<3:1 | <<3:1 | <<3:1 | <<3:1 | <<3:1 | >>3:1 | >>3:1 | >>3:1 |
| ZG184 | <<3:1 | <<3:1 | <<3:1 | <<3:1 | <<3:1 | <<3:1 | <<3:1 | <<3:1 | <<3:1 | >>3:1 | >>3:1 | >>3:1 |
| ZG185 | 8.0:1 | 7.2:1 | 5.9:1 | 7.3:1 | 9.8:1 | 6.0:1 | 4.8:1 | 3.3:1 | 1.7:1 | 7.9:1 | 20.4:1 | 7.0:1 |
| ZG186 | 5.2:1 | 3.5:1 | 2.0:1 | 7.6:1 | 6.6:1 | 5.0:1 | 9.7:1 | 5.9:1 | 1.5:1 | 4.4:1 | 5.0:1 | 7.8:1 |
| ZG187 | 10.8:1 | 4.7:1 | 3.7:1 | 9.9:1 | 14.7:1 | 4.6:1 | 6.8:1 | 5.1:1 | 2.0:1 | >>3:1 | >>3:1 | >>3:1 |
| ZG188 | <<3:1 | <<3:1 | <<3:1 | 9.4:1 | 5.0:1 | 3.8:1 | <<3:1 | <<3:1 | <<3:1 | <<3:1 | <<3:1 | <<3:1 |
| ZG189 | <<3:1 | <<3:1 | <<3:1 | 4.9:1 | 6.8:1 | 5.1:1 | <<3:1 | <<3:1 | <<3:1 | <<3:1 | <<3:1 | <<3:1 |
| ZG190 | <<3:1 | <<3:1 | <<3:1 | 7.0:1 | 4.6:1 | 3.3:1 | <<3:1 | <<3:1 | <<3:1 | <<3:1 | <<3:1 | <<3:1 |
| ZG194 | <<3:1 | <<3:1 | <<3:1 | <<3:1 | <<3:1 | <<3:1 | 20.1:1 | 5.8:1 | 1.3:1 | <<3:1 | <<3:1 | <<3:1 |
| ZG195 | <<3:1 | <<3:1 | <<3:1 | <<3:1 | <<3:1 | <<3:1 | 9.0:1 | 3.9:1 | 3.1:1 | <<3:1 | <<3:1 | <<3:1 |
| ZG196 | <<3:1 | <<3:1 | <<3:1 | <<3:1 | <<3:1 | <<3:1 | 7.6:1 | 4.6:1 | 1.2:1 | <<3:1 | <<3:1 | <<3:1 |
| ZG197 | 5.0:1 | 4.8:1 | 1.9:1 | <<3:1 | <<3:1 | <<3:1 | <<3:1 | <<3:1 | <<3:1 | 1.5:1 | 3.8:1 | 0.9:1 |
| ZG198 | 9.1:1 | 4.3:1 | 4.8:1 | <<3:1 | <<3:1 | <<3:1 | <<3:1 | <<3:1 | <<3:1 | 8.8:1 | 5.9:1 | 11.8:1 |
| ZG199 | 34.8:1 | 6.0:1 | 7.3:1 | <<3:1 | <<3:1 | <<3:1 | <<3:1 | <<3:1 | <<3:1 | 9.4:1 | 11.7:1 | 10.7:1 |

| | YISLIYTNYEAGKDDYVK | | | VSAMYSSSSCKLPSLSPVAR | | | EHCSACGPLSR | | | EWSTFAVGPGHCLQLNDR | | |
|---|---|---|---|---|---|---|---|---|---|---|---|---|
| Replicate | y8 - 895.4520+ | b2 - 277.1547+ | b3 - 364.1867+ | y8 - 826.4781+ | y5 - 529.3093+ | b3 - 258.1448+ | y9 - 1007.4397+ | y8 - 847.4091+ | b3 - 427.1394+ | y5 - 645.3315+ | y4 - 517.2729+ | b6 - 722.3144+ |
| ZG182 | 5.3:1 | 54.2:1 | 13.5:1 | <<3:1 | <<3:1 | <<3:1 | 18.2:1 | 9.9:1 | 2.9:1 | <<3:1 | <<3:1 | <<3:1 |
| ZG183 | 4.4:1 | 56.8:1 | 13.3:1 | <<3:1 | <<3:1 | <<3:1 | 56.0:1 | 13.9:1 | 8.9:1 | <<3:1 | <<3:1 | <<3:1 |
| ZG184 | 4.4:1 | 33.0:1 | 15.4:1 | <<3:1 | <<3:1 | <<3:1 | 33.1:1 | 14.9:1 | 6.5:1 | <<3:1 | <<3:1 | <<3:1 |
| ZG185 | 1.7:1 | 13.9:1 | 7.9:1 | <<3:1 | <<3:1 | <<3:1 | 31.1:1 | 28.8:1 | 6.6:1 | <<3:1 | <<3:1 | <<3:1 |
| ZG186 | 0.8:1 | 22.2:1 | 4.9:1 | <<3:1 | <<3:1 | <<3:1 | 24.5:1 | 19.6:1 | 9.2:1 | <<3:1 | <<3:1 | <<3:1 |
| ZG187 | 0.5:1 | 24.2:1 | 10.0:1 | <<3:1 | <<3:1 | <<3:1 | 42.7:1 | 41.6:1 | 7.1:1 | <<3:1 | <<3:1 | <<3:1 |
| ZG188 | 0.8:1 | 17.9:1 | 3.6:1 | <<3:1 | <<3:1 | <<3:1 | 23.5:1 | 23.5:1 | 7.1:1 | <<3:1 | <<3:1 | <<3:1 |
| ZG189 | 3.1:1 | 15.0:1 | 6.6:1 | <<3:1 | <<3:1 | <<3:1 | 30.0:1 | 20.6:1 | 4.5:1 | <<3:1 | <<3:1 | <<3:1 |
| ZG190 | 0.7:1 | 19.2:1 | 10.3:1 | <<3:1 | <<3:1 | <<3:1 | 75.3:1 | 75.0:1 | 5.6:1 | <<3:1 | <<3:1 | <<3:1 |
| ZG194 | <<3:1 | <<3:1 | <<3:1 | <<3:1 | <<3:1 | <<3:1 | 14.9:1 | 15.9:1 | 2.1:1 | <<3:1 | <<3:1 | <<3:1 |
| ZG195 | <<3:1 | <<3:1 | <<3:1 | <<3:1 | <<3:1 | <<3:1 | 10.3:1 | 13.8:1 | 1.9:1 | <<3:1 | <<3:1 | <<3:1 |
| ZG196 | <<3:1 | <<3:1 | <<3:1 | <<3:1 | <<3:1 | <<3:1 | 9.3:1 | 3.7:1 | 2.1:1 | <<3:1 | <<3:1 | <<3:1 |
| ZG197 | 0.9:1 | 6.9:1 | 2.3:1 | <<3:1 | <<3:1 | <<3:1 | 16.5:1 | 12.1:1 | 7.4:1 | <<3:1 | <<3:1 | <<3:1 |
| ZG198 | 0.8:1 | 5.2:1 | 2.1:1 | <<3:1 | <<3:1 | <<3:1 | 26.5:1 | 18.4:1 | 9.5:1 | <<3:1 | <<3:1 | <<3:1 |
| ZG199 | 0.5:1 | 10.8:1 | 2.9:1 | <<3:1 | <<3:1 | <<3:1 | 43.8:1 | 26.6:1 | 8.9:1 | <<3:1 | <<3:1 | <<3:1 |

| | YVSLIYTNYEAGKDDYVK | | | VSAMYSSSPCKLPSLSPVAR | | | EHCSACGPLSQLLVK | | | EWSTFAVGPGHCLQLHDR | | |
|---|---|---|---|---|---|---|---|---|---|---|---|---|
| Replicate | y12 - 1402.6485+ | b2 - 263.1390+ | b3 - 350.1710+ | y8 - 826.4781+ | y5 - 529.3093+ | b3 - 258.1448+ | y13 - 1432.7287+ | y9 - 954.5982+ | b3 - 427.1394+ | y11 - 1289.6168+ | y3 - 427.2048+ | b6 - 722.3144+ |
| ZG182 | <<3:1 | <<3:1 | <<3:1 | <<3:1 | <<3:1 | <<3:1 | <<3:1 | <<3:1 | <<3:1 | <<3:1 | <<3:1 | <<3:1 |
| ZG183 | <<3:1 | <<3:1 | <<3:1 | <<3:1 | <<3:1 | <<3:1 | <<3:1 | <<3:1 | <<3:1 | <<3:1 | <<3:1 | <<3:1 |
| ZG184 | <<3:1 | <<3:1 | <<3:1 | <<3:1 | <<3:1 | <<3:1 | <<3:1 | <<3:1 | <<3:1 | <<3:1 | <<3:1 | <<3:1 |
| ZG185 | <<3:1 | <<3:1 | <<3:1 | <<3:1 | <<3:1 | <<3:1 | <<3:1 | <<3:1 | <<3:1 | <<3:1 | <<3:1 | <<3:1 |
| ZG186 | <<3:1 | <<3:1 | <<3:1 | <<3:1 | <<3:1 | <<3:1 | <<3:1 | <<3:1 | <<3:1 | <<3:1 | <<3:1 | <<3:1 |
| ZG187 | <<3:1 | <<3:1 | <<3:1 | <<3:1 | <<3:1 | <<3:1 | <<3:1 | <<3:1 | <<3:1 | <<3:1 | <<3:1 | <<3:1 |
| ZG188 | <<3:1 | <<3:1 | <<3:1 | <<3:1 | <<3:1 | <<3:1 | <<3:1 | <<3:1 | <<3:1 | <<3:1 | <<3:1 | <<3:1 |
| ZG189 | <<3:1 | <<3:1 | <<3:1 | <<3:1 | <<3:1 | <<3:1 | <<3:1 | <<3:1 | <<3:1 | <<3:1 | <<3:1 | <<3:1 |
| ZG190 | <<3:1 | <<3:1 | <<3:1 | <<3:1 | <<3:1 | <<3:1 | <<3:1 | <<3:1 | <<3:1 | <<3:1 | <<3:1 | <<3:1 |
| ZG194 | <<3:1 | <<3:1 | <<3:1 | <<3:1 | <<3:1 | <<3:1 | <<3:1 | <<3:1 | <<3:1 | <<3:1 | <<3:1 | <<3:1 |
| ZG195 | <<3:1 | <<3:1 | <<3:1 | <<3:1 | <<3:1 | <<3:1 | <<3:1 | <<3:1 | <<3:1 | <<3:1 | <<3:1 | <<3:1 |
| ZG196 | <<3:1 | <<3:1 | <<3:1 | <<3:1 | <<3:1 | <<3:1 | <<3:1 | <<3:1 | <<3:1 | <<3:1 | <<3:1 | <<3:1 |
| ZG197 | <<3:1 | <<3:1 | <<3:1 | <<3:1 | <<3:1 | <<3:1 | <<3:1 | <<3:1 | <<3:1 | <<3:1 | <<3:1 | <<3:1 |
| ZG198 | <<3:1 | <<3:1 | <<3:1 | <<3:1 | <<3:1 | <<3:1 | <<3:1 | <<3:1 | <<3:1 | <<3:1 | <<3:1 | <<3:1 |
| ZG199 | <<3:1 | <<3:1 | <<3:1 | <<3:1 | <<3:1 | <<3:1 | <<3:1 | <<3:1 | <<3:1 | <<3:1 | <<3:1 | <<3:1 |

| | GVALSNVIHK | | | DLNMDCMVAEIK | | | GAFLYEPCGVSTPVLSTGVLR | | | ARPLEQAVAAIVCTFQEYAGR | | |
|---|---|---|---|---|---|---|---|---|---|---|---|---|
| Replicate | y6 - 697.3991+ | y3 - 397.2558+ | b3 - 228.1343+ | y8 - 965.4431+ | y6 - 690.3855+ | b3 - 343.1612+ | y9 - 941.5778+ | b6 - 681.3243+ | b10 - 1094.4975+ | y13 - 1485.7155+ | b2 - 270.1561+ | b11 - 1162.6579+ |
| ZG182 | 1.5:1 | 1.7:1 | 1.8:1 | 1.7:1 | 0.5:1 | 4.1:1 | 1.9:1 | 0.1:1 | 2.9:1 | >>3:1 | >>3:1 | >>3:1 |
| ZG183 | 2.8:1 | 1.0:1 | 3.3:1 | <<3:1 | <<3:1 | <<3:1 | 3.4:1 | 0.1:1 | 2.5:1 | >>3:1 | >>3:1 | >>3:1 |
| ZG184 | 5.2:1 | 3.1:1 | 5.7:1 | <<3:1 | <<3:1 | <<3:1 | 1.2:1 | 0.3:1 | 1.4:1 | >>3:1 | >>3:1 | >>3:1 |
| ZG185 | 4.1:1 | 2.7:1 | 2.3:1 | <<3:1 | <<3:1 | <<3:1 | >>3:1 | >>3:1 | >>3:1 | >>3:1 | >>3:1 | >>3:1 |
| ZG186 | 5.0:1 | 1.9:1 | 4.7:1 | <<3:1 | <<3:1 | <<3:1 | >>3:1 | >>3:1 | >>3:1 | >>3:1 | >>3:1 | >>3:1 |
| ZG187 | 4.8:1 | 1.4:1 | 6.7:1 | <<3:1 | <<3:1 | <<3:1 | >>3:1 | >>3:1 | >>3:1 | >>3:1 | >>3:1 | >>3:1 |
| ZG188 | 1.5:1 | 2.5:1 | 3.0:1 | <<3:1 | <<3:1 | <<3:1 | 12.1:1 | 37.5:1 | 2.5:1 | >>3:1 | >>3:1 | >>3:1 |
| ZG189 | 3.0:1 | 1.5:1 | 3.4:1 | <<3:1 | <<3:1 | <<3:1 | 10.6:1 | 37.8:1 | 5.3:1 | >>3:1 | >>3:1 | >>3:1 |
| ZG190 | 1.9:1 | 2.3:1 | 5.1:1 | <<3:1 | <<3:1 | <<3:1 | 13.5:1 | 60.7:1 | 4.4:1 | >>3:1 | >>3:1 | >>3:1 |
| ZG194 | 1.4:1 | 0.2:1 | 2.2:1 | <<3:1 | <<3:1 | <<3:1 | <<3:1 | <<3:1 | <<3:1 | >>3:1 | >>3:1 | >>3:1 |
| ZG195 | 1.0:1 | 1.0:1 | 1.6:1 | <<3:1 | <<3:1 | <<3:1 | <<3:1 | <<3:1 | <<3:1 | >>3:1 | >>3:1 | >>3:1 |
| ZG196 | 0.8:1 | 2.5:1 | 2.7:1 | <<3:1 | <<3:1 | <<3:1 | <<3:1 | <<3:1 | <<3:1 | >>3:1 | >>3:1 | >>3:1 |
| ZG197 | 4.3:1 | 5.9:1 | 6.6:1 | <<3:1 | <<3:1 | <<3:1 | <<3:1 | <<3:1 | <<3:1 | <<3:1 | <<3:1 | <<3:1 |
| ZG198 | 1.8:1 | 0.9:1 | 5.2:1 | 6.4:1 | 1.4:1 | 7.9:1 | <<3:1 | <<3:1 | <<3:1 | <<3:1 | <<3:1 | <<3:1 |
| ZG199 | 5.5:1 | 0.4:1 | 2.1:1 | 13.9:1 | 2.0:1 | 15.2:1 | <<3:1 | <<3:1 | <<3:1 | <<3:1 | <<3:1 | <<3:1 |

| | GVALSNVVHK | | | DLNMDCIVAEIK | | | GAFLYDPCGVSTPVLSTGVLR | | | AKPLEQAVAAIVCTFQEYAGR | | |
|---|---|---|---|---|---|---|---|---|---|---|---|---|
| Replicate | y8 - 867.5047+ | y7 - 796.4676+ | y6 - 683.3835+ | y8 - 947.4866+ | y6 - 672.4291+ | b3 - 343.1612+ | y9 - 941.5778+ | b6 - 667.3086+ | b10 - 1080.4819+ | y13 - 1485.7155+ | b2 - 242.1499+ | b11 - 1134.6517+ |
| ZG182 | 21.9:1 | 5.8:1 | 7.2:1 | >>3:1 | >>3:1 | >>3:1 | <<3:1 | <<3:1 | <<3:1 | <<3:1 | <<3:1 | <<3:1 |
| ZG183 | 38.6:1 | 1.8:1 | 4.6:1 | >>3:1 | >>3:1 | >>3:1 | <<3:1 | <<3:1 | <<3:1 | <<3:1 | <<3:1 | <<3:1 |
| ZG184 | 25.3:1 | 4.3:1 | 7.0:1 | >>3:1 | >>3:1 | >>3:1 | <<3:1 | <<3:1 | <<3:1 | <<3:1 | <<3:1 | <<3:1 |
| ZG185 | 21.1:1 | 3.6:1 | 7.8:1 | >>3:1 | >>3:1 | >>3:1 | <<3:1 | <<3:1 | <<3:1 | 24.7:1 | 7.9:1 | 10.4:1 |
| ZG186 | 19.7:1 | 3.7:1 | 6.7:1 | >>3:1 | >>3:1 | >>3:1 | <<3:1 | <<3:1 | <<3:1 | 22.7:1 | 6.3:1 | 17.8:1 |
| ZG187 | 19.5:1 | 4.5:1 | 9.0:1 | >>3:1 | >>3:1 | >>3:1 | <<3:1 | <<3:1 | <<3:1 | 64.5:1 | 16.5:1 | 14.3:1 |
| ZG188 | 12.0:1 | 4.4:1 | 6.7:1 | >>3:1 | >>3:1 | >>3:1 | <<3:1 | <<3:1 | <<3:1 | <<3:1 | <<3:1 | <<3:1 |
| ZG189 | 27.1:1 | 4.3:1 | 4.1:1 | >>3:1 | >>3:1 | >>3:1 | <<3:1 | <<3:1 | <<3:1 | <<3:1 | <<3:1 | <<3:1 |
| ZG190 | 4.9:1 | 3.4:1 | 5.1:1 | >>3:1 | >>3:1 | >>3:1 | <<3:1 | <<3:1 | <<3:1 | <<3:1 | <<3:1 | <<3:1 |
| ZG194 | 6.9:1 | 3.2:1 | 4.1:1 | <<3:1 | <<3:1 | <<3:1 | 15.8:1 | 21.1:1 | 2.7:1 | 14.6:1 | 2.0:1 | 5.7:1 |
| ZG195 | 6.4:1 | 1.9:1 | 2.2:1 | <<3:1 | <<3:1 | <<3:1 | 6.9:1 | 6.4:1 | 2.7:1 | 16.4:1 | 4.7:1 | 7.9:1 |
| ZG196 | 12.1:1 | 2.7:1 | 5.8:1 | <<3:1 | <<3:1 | <<3:1 | 14.0:1 | 19.8:1 | 3.2:1 | 9.9:1 | 1.6:1 | 3.1:1 |
| ZG197 | <<3:1 | <<3:1 | <<3:1 | <<3:1 | <<3:1 | <<3:1 | 3.0:1 | 2.7:1 | 1.0:1 | <<3:1 | <<3:1 | <<3:1 |
| ZG198 | <<3:1 | <<3:1 | <<3:1 | <<3:1 | <<3:1 | <<3:1 | 4.4:1 | 7.5:1 | 1.9:1 | <<3:1 | <<3:1 | <<3:1 |
| ZG199 | <<3:1 | <<3:1 | <<3:1 | <<3:1 | <<3:1 | <<3:1 | 15.5:1 | 12.6:1 | 2.9:1 | <<3:1 | <<3:1 | <<3:1 |

Figure S5. **MRM signal to noise ratios**. Signal to noise ratios are reported as the intensity of the signal divided by the amplitude of the noise. Values that visually appear much higher than 3:1 are reported as >>3:1 and values that are visually much lower than 3:1 are reported as <<3:1. ZG182-184 = E1, ZG185-187 = E2, ZG188-190 = E3, ZG194-196 = A1, ZG197-199 = E2.

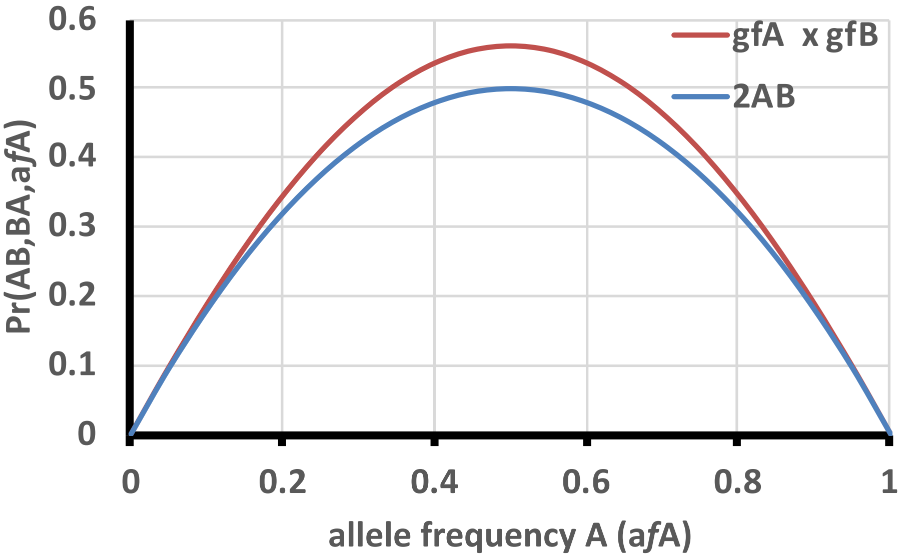


Figure S6. **Calculation of Heterozygote Frequency Using Product of Genotype Frequencies.** The MCMC model of estimated random match probabilities approximates the calculation of heterozygote probability (2AB) by substituting the product of respective genotype frequencies of each alleles (g*f*A x g*f*B). The maximum difference between the two estimates is Pr = 0.065, resulting in slightly higher probabilities.
